# The structure of insulin granule core determines secretory capacity being reduced in type-2 diabetes

**DOI:** 10.1101/2022.02.22.481455

**Authors:** Mohammad Barghouth, Xiaoping Jiang, Mototsugu Nagao, Ning Chen, Daowei Yang, Yingying Ye, Cheng Luan, Maria F. Gomez, Anna M. Blom, Claes B Wollheim, Lena Eliasson, Erik Renström, Enming Zhang

## Abstract

Exocytosis in excitable cells is essential for their physiological functions. Although the exocytotic machinery controlling cellular secretion has been well investigated, the function of the vesicular cargo, i.e. secretory granular content remains obscure. Here we combine dSTORM imaging and single-domain insulin antibody, to dissect the *in situ* structure of insulin granule cores (IGCs) at nano level. We demonstrate that the size and shape of the IGCs can be regulated by the juxta-granular molecules Nucleobindin-2 and Enolase-1, that further contribute to the stimulated insulin secretion. IGCs located at the plasma membrane are larger than those in the cytosol. The IGCs size is decreased by ∼20% after glucose stimulation due to the release of the peripheral part of IGCs through incomplete granule fusion. Importantly, the reduction of the IGCs size is also observed in non-stimulatory pancreatic β-cells from diabetic *db/db* mice, Akita (*Ins2^+/-^*) mice and human Type-2 diabetic donors, in accordance with impaired secretion. These findings overall highlight the structure of exocytotic insulin cores as a novel modality amenable to targeting in the stimulated exocytosis in β-cells with impaired insulin secretion.

## Introduction

Exocytosis is the essential function of excitable and other secretory cells e.g. neurons, chromaffin cells and pancreatic β−cells. Defective exocytosis underlies disorders and diseases, including neurodegenerative diseases such as Alzheimer’s disease, as well as type-2 diabetes (T2D). During the past decades, stimulated exocytosis has been thoroughly investigated, establishing the mechanisms through which the secretory machinery determines granule trafficking, docking, priming, fusion and refilling (Eliasson et al., 2008; Gandasi et al., 2018; Sudhof and Rothman, 2009). Exocytosis comprises granule tethering to the components of the cytoskeleton and calcium-triggered SNARE (soluble N-ethylmaleimide-sensitive factor attachment protein receptor) complex mediated priming and fusion(Sudhof and Rothman, 2009). In contrast to the molecular control of the triggering of exocytosis, how cargo components and structures contribute to the capacity of secretion remains to be elucidated.

Impaired insulin secretion from the pancreatic β-cells is the main cause of T2D (Lyssenko et al., 2008; Thurmond and Gaisano, 2020; Zhang et al., 2019), which is the most common endocrine disorder worldwide (Ligthart et al., 2016). Insulin is stored in dense-core granules in the β-cell, the release of which is evoked by a series of subcellular physiological events, including glucose transport and metabolization, ATP production in the mitochondria, closure of ATP-sensitive K^+^ channels, plasma membrane depolarization, and finally Ca^2+^ influx that triggers insulin granule fusion (Del Prato et al., 2002; Eliasson *et al*., 2008; Thurmond and Gaisano, 2020). The insulin granule contains ∼150 protein species, such as either granule membrane proteins or cargo contents (Hutton et al., 1982; Suckale and Solimena, 2010). Predominant granule transmembrane proteins include synaptotagmins and VAMP2 (Regazzi et al., 1996). The latter is part of the SNARE complex, together with the plasma membrane proteins syntaxin 1 and SNAP25, controlling insulin secretion by regulating granule exocytosis (Andersson et al., 2012; Gandasi *et al*., 2018; Ostenson et al., 2006; Rorsman and Renstrom, 2003; Thurmond and Gaisano, 2020). Inside the granule, insulin accounts for 50-60% of cargo protein content (Hutton *et al*., 1982). The remainder comprises proteins such as islet amyloid polypeptide (IAPP, also known as amylin) that is co-secreted with insulin at ∼1:3 ratio(Ogawa et al., 1990); GABA, which is estimated to account for 15% of total protein co-secreted with insulin; and IGF2(Cornu et al., 2009), granins and granin-derived peptides, as well as prohormone convertases (Brunner et al., 2007), which play essential roles in insulin granule maturation and ensure processing of proinsulin to insulin. The prohormone convertases and soluble molecules in the granules, such as ATP, H^+^, Ca^2+^ and Zn^2+^ (necessary for the insulin crystal formation), are also crucial for maintaining granule function and release competence (Davidson et al., 1988; Lemaire et al., 2009; Orci et al., 1986). These cargo proteins, including insulin, form an ultrastructural electron-dense complex core detectable by transmission electron microscopy (TEM) (Hoboth et al., 2015; Nam et al., 2014). However, the pure insulin core, which excludes other cargo proteins, has not been defined by existing imaging methods and its putative function beyond simple storage form remains to be elucidated.

Direct stochastic optical reconstruction microscopy (dSTORM) has marked advantages with high-precision localization of single molecules beyond the optical resolution limit (Heilemann et al., 2008; Rust et al., 2006). Since detection is performed with standard fluorescent dyes, this method can be used in most biological samples and tissues to distinguish specific proteins, potentially contributing to pathological diagnosis (Hu et al., 2016; Mengistu et al., 2017). Several factors can affect the resolution of dSTORM imaging, of which the most important is the probe size(Dempsey et al., 2011). The probe complex, including primary, secondary antibodies (∼10 nm each) and fluorescent dye, is more than 20 nm in total size, which limits further increasing the resolution of detection in super-resolution imaging (Ries et al., 2012). Recently, a single domain (SD) antibody against insulin was developed with a remarkably smaller size (<2 nm)(Ries *et al*., 2012; Rothbauer et al., 2006), which provides the possibility to improve the resolution of dSTORM images. Further, the SD antibody can pass small holes, e.g., the exocytotic granule’s fusion pore, which averages a few nanometers. To this end, we developed a state-of-art method to image IGCs by using the anti-insulin SD antibody in combination with dSTORM. Using this approach, we demonstrated that the pure insulin core in the granule is dynamically changing during glucose stimulation and can be regulated by either granular proteins or extracellular stimuli. Furthermore, the size reduction is related to incomplete granule fusion and characterizes defective exocytosis in beta cells in T2D donors.

## Results

### Detection of *in situ* IGCs using SD antibody probed by dSTORM imaging

The recently developed SD antibody (1∼2 nm, 10-15 kDa), is much smaller than common IgG antibodies (10 nm, 150-160 kDa) and therefore favourably for nano-scale imaging (Ries *et al*., 2012). To take advantage of this size difference in structure imaging of insulin granule cores (IGCs), we performed the super-resolution dSTORM imaging with camelid SD antibodies against insulin in INS-1 832-13 cells. Although the cores can be detected with the normal IgG antibody by the dSTORM, the staining pattern of the insulin cores appeared scattered and irregular, with potentially unspecific dots (Figure 1A). In comparison, the SD antibody stained the insulin core in a condensed fashion with smooth and clear edges (Figure 1B, Figure S1A and B). We next measured the area, average diameter and perimeter of the insulin core. Interestingly, all insulin core parameters in SD antibody stainings were larger than those seen with IgG antibodies (Figure 1C and Figure S1C-E). With SD antibodies, the average core area and the perimeter was 14013±575 nm^2^ and 347±8.6 nm, in comparison with the IgG antibody, respectively 8149±306 nm^2^ and 254±5.7 nm. The average diameters of insulin cores binned into size groups showed that the population of insulin cores >100 nm was increased 60% by SD antibody (Figure S1F), while the population of the small cores (40-60 nm) was increased by IgG antibody staining. In total, the average diameter of insulin cores stained by SD antibody was 118±23 nm and by IgG antibodies is 83±13 nm (Figure 1D). In contrast, with conventional imaging the average diameter of IGCs did not differ significantly between the two types of antibody (Figure 1E).

**Figure 1.**
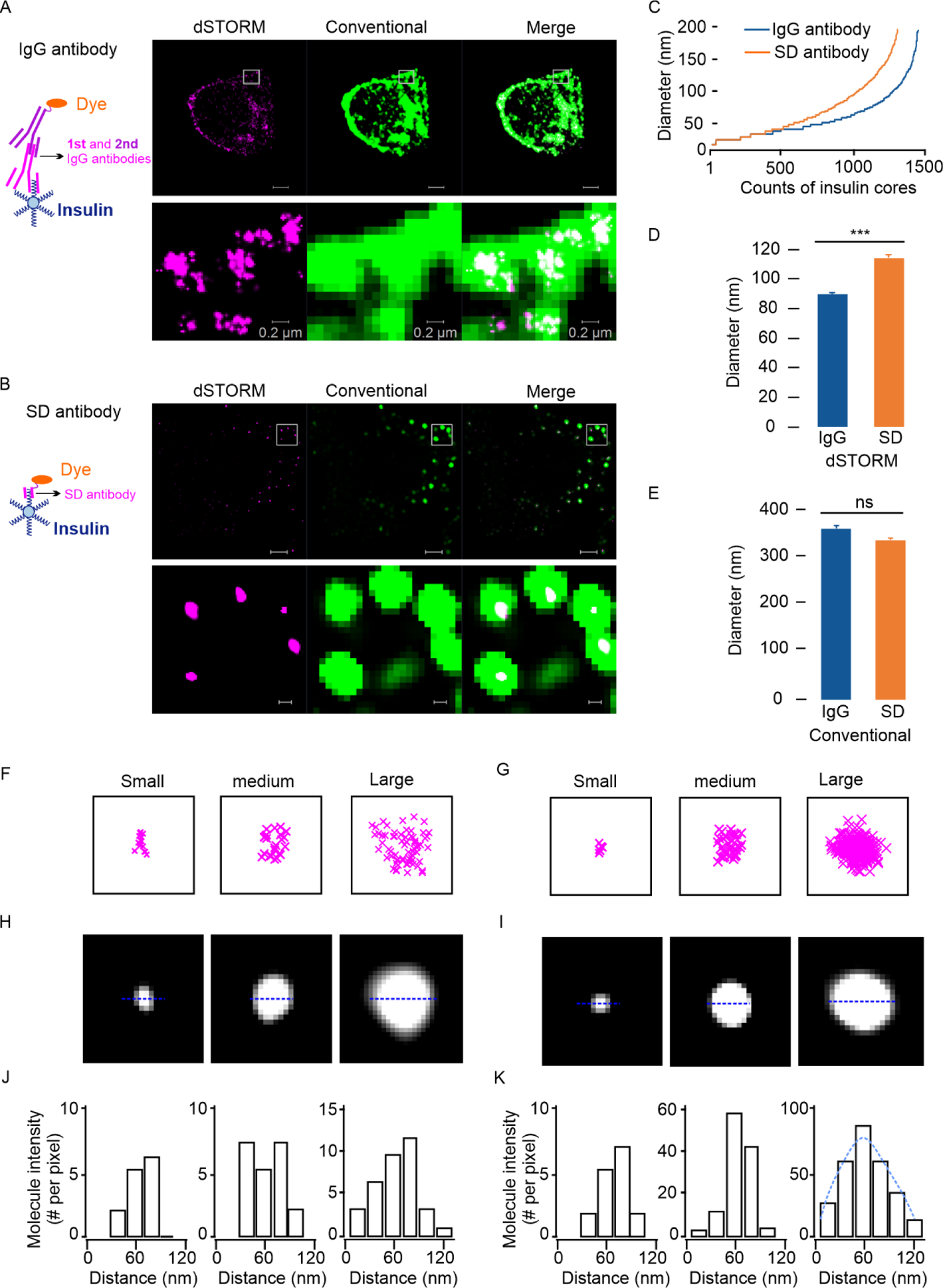
Comparison of insulin core sizes as detected by IgG and single domain (SD) insulin antibodies. (A) Insulin core size detection in insulin-secreting Ins-1(832/13) cells by an IgG primary insulin antibody plus the second antibody conjugated with Alexa 647 (left) with representative dSTORM images detected (right). Magnified images are cut from the squared regions indicated in the upper images (right bottom). (B) Same as in (A), but detection is done using single domain antibodies conjugated with Alexa 647. (C) Diameter of insulin cores using IgG or SD antibodies detected by dSTORM imaging. The data was collected from 1280 granules by IgG and 1450 granules by SD in 18 cells. (D) The average diameters of insulin cores were larger when assessed using SD antibodies than by IgG antibodies in dSTORM images, but no difference was seen in conventional images (E). (F) Insulin molecule density in small (left, ∼60 nm), medium (∼90 nm) or large (right, ∼120 nm) insulin cores measured with IgG insulin antibodies or (G) with SD insulin antibodies. (H) Selected insulin cores of small, medium and large sizes were detected by IgG insulin antibodies. (I) same as in (H) but detected by SD insulin antibodies. (J) The distribution of molecular intensity (the insulin molecules per pixel measured following the equatorial section of the insulin cores) when measured in insulin core of small, medium and large sizes with IgG insulin antibodies and (K) with SD insulin antibodies. Data presented as means ± SEM and collected from 18 cells in three independent experiments. * p<0.05; ns, no significance.

### SD antibody probes penetrate predominantly into insulin cores

To evaluate whether the larger IGCs detected by SD antibodies reflects an according amount of insulin molecules, we next analyzed photon number and their locations in insulin cores. The analysis showed that the IGCs can be classified into three subpopulations: small (∼60 nm), medium (∼90 nm) and large (∼120 nm) diameter insulin cores (Figure 1F-I). The photon density distribution (calculated by photon number per pixel) over an insulin core appeared irregular when staining with IgG antibodies (Figure 1J), indicating that the insulin within the cores was unsaturated with regards to binding insulin antibodies. In contrast, the SD antibody-stained IGCs appeared to display a more Gaussian distribution, particularly in the large subgroup of insulin cores (Figure 1K). The blinking photons reflecting the bound SD antibodies were much higher in the large size cores, in which the average photon numbers were 2585±210 vs 414±28 in IgG-stained cores (Figure S1G, H). overall, these results demonstrate that insulin molecules within IGCs can be easily detected by the SD antibodies compared to IgG antibodies.

### IGCs with larger size locate near the plasma membrane

To estimate the homogeneity of the spatial distribution of IGCs, we used two acquisition modes of the dSTORM imaging to detect the size of all insulin cores in intact cells: 1) epifluorescent mode (EPI) used to detect most insulin cores in the cytosol; 2) total internal reflection (TIRF) mode used to visualize insulin cores near the plasma membrane (< 150 nm) (Figure S2A). Under TIRF mode, the proportion of large-size (>80 nm) insulin cores accounted for >80% of insulin cores, while under EPI mode, <30% of insulin cores had a diameter larger than 80 nm (Figure 2A-C). Accordingly, the area of insulin cores was ∼25000 nm^2^ by the TIRF mode, whereas, in the EPI mode, it was estimated to be only ∼8000 nm^2^ (Figure S2B). These large IGCs were preferentially located close to the plasma membrane. Next, the homogeneity of IGCs distribution was estimated by calculating the correlation between area and perimeter (Figure S2C). Compared to the EPI mode, the fitting curve of the plots under TIRF mode resulted in a linear distribution in the span of IGCs with large size, indicating that insulin cores tended to shift from elliptical shape to round when approaching the plasma membrane. These analyses showed that the shape of IGCs changes dynamically according to their position relative to the plasma membrane. This phenomenon may be linked to the exocytotic process.

**Figure 2.**
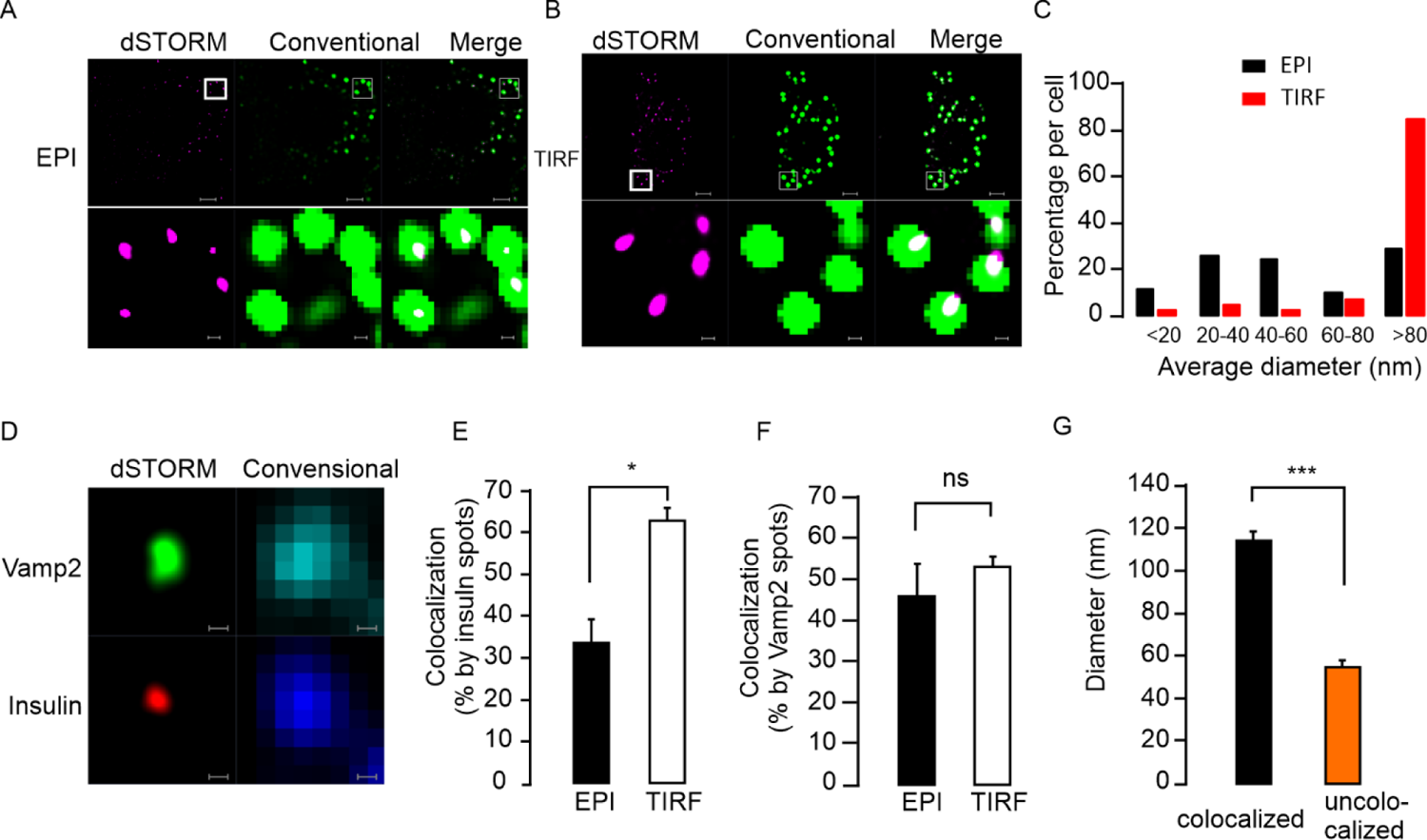
Large insulin cores preferentially locate near the plasma membrane. (A) dSTORM images of insulin cores using SD antibodies in the epifluorescence (EPI) mode or (B) Total internal reflection (TIRF) mode. The magnifications (lower panels) represent the squares indicated in the upper images. (C) Distribution of insulin core diameters as measured in EPI or TIRF mode. (D) A single insulin core colocalized with exocytotic marker Vamp2 visualized by two-colour dSTORM imaging. (E) Fraction of insulin cores colocalized with Vamp2 to total insulin cores in EPI or TIRF mode. (F) same as in (E), but when assessing the fraction of Vamp2 clusters colocalized with insulin cores to total number of Vamp2 clusters. (G) Comparison of average insulin core diameters when colocalized, or not with Vamp2. Data presented are means ± SEM and collected from 21 INS-1(832/13) cells under each condition from 3 independent experiments. Significance test performed with single factor ANOVA test. * p<0.05; ***, p<0.001; ns, no significance.

### The exocytotic IGCs are detected under TIRF mode

Exocytotic granules are large-dense core vesicles, which are supposed to be tethered and docked on the plasma membrane waiting for triggering signals in order to release their contents (Wang and Thurmond, 2009). To prove that the large IGCs are located in the exocytotic granules, we co-stained insulin with Vamp2, a membrane marker of exocytotic granules (Regazzi *et al*., 1996) by two-colour dSTORM imaging (Figure 2D). Fortunately, using an IgG Vamp2 antibody, we could clearly acquire the dSTORM images of Vamp2, likely because of their scattered distribution in the granule membrane (Figure S2D and F). The colocalization of the insulin cores and VAMP2 was preferentially observed juxta-plasma membrane rather than in the cytosol (Figure S2E and G). Statistically, the IGCs that colocalized with Vamp2 were 63±3% at the plasma membrane concomitant with 34±6% in the cytosol (Figure 2E). However, the percentage of Vamp2, which colocalized with insulin cores, did not differ between cytosol and plasma membrane (Figure 2F), indicating that Vamp2 has no preferable selection for its physical locations in the cells. However, the IGCs that colocalized with Vamp2 were larger than that not colocalizing (Figure 2G). These results suggest that exocytotic IGCs with large sizes located at the plasma membrane are ready for secretion.

### Granule proteins participate in the regulation of IGC size and proinsulin/insulin composition

Proteomics analysis showed that 51 proteins reside in Vamp2 marked exocytotic granules(Hickey et al., 2009). Among these, we have selected 11 candidates (CPE, Znt8, ChgA, Eno 1, ATP6V0a1, ATP6V1a, Ctsd, Pdia3, Pcsk2, Nucb2) to assess their roles in the regulation of IGC’s morphology. After silencing the proteins individually in INS-1 832-13 cells (Figure S3A), the size of the IGCs located in the cytosol or juxta-plasma membrane were detected by dSTORM imaging under EPI or TIRF mode, respectively. Of note, silencing Pcsk2 (Prohormone Convertase 2) or Nucb2 (Nucleobindin 2) significantly increased the cytosolic core sizes under EPI mode, while silencing CPE (Carboxypeptidase E), PDIA3 (Protein Disulfide Isomerase-Associated 3), or Ctsd (Cathepsin D) enlarged the juxta-membrane cores visualized under TIRF mode (Figure 3A and 3B). Besides the size, the core shape is indicated by the polarizability index (Figure S3B), which shows the shift of insulin cores from elliptical to round shape. The index of polarizability is 1 for the absolute round IGCs, while the value is approximately 0 in the long elliptical shape. The cytosolic IGCs detected by EPI mode yield 0.47±0.06 with a slight oval shape. The polarizability index of surface insulin cores was 0.58±0.05 and slightly rounder than cytosolic insulin cores (Figure 3C and D). Especially silencing Eno1 caused rounder shape than control significantly under TIRF mode (0.70±0.03 vs 0.58±0.05) (Figure 3D). These analyses showed that certain granule proteins such as Nucb2 and Eno-1 are required for maintaining the IGC structure and shape.

**Figure 3.**
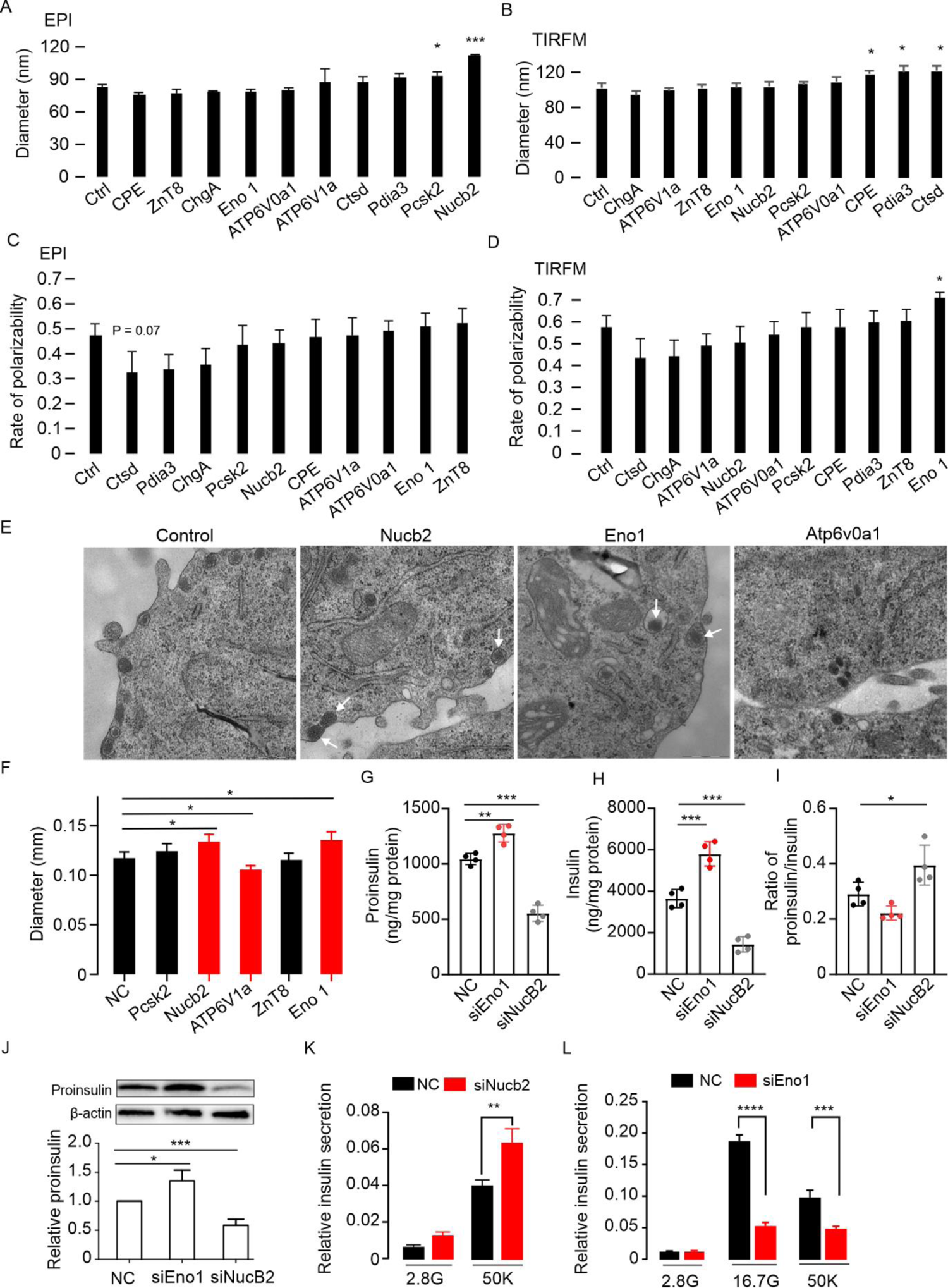
Insulin granular proteins are involved in the regulation of insulin core size. (A) Diameter of insulin cores detected by dSTORM imaging under EPI (A) or TIRF mode (B) after 48 h knock-down of the granular proteins. The rate of the polarizability of the insulin core shape was detected under the same conditions as in (A) under EPI (C) or TIRF mode (D). (E) Representative TEM images of insulin granules after 48 h knock-down of the granular proteins. The arrows indicate the enlarged insulin cores in the Eno 1 or Nucleobindin 2 KD INS-1(832/13) cells. (F) Diameter of insulin cores detected by TEM imaging in the KD cells. Proinsulin (G), insulin (H) and ration of proinsulin/insulin (I) were measured in the Eno-1 and Nucb2 silencing INS-1 cells. (J) The proinsulin in the cells was measured by western blot. (K) The insulin secretion increased in the Nucleobindin 2 KD cells after 50 mM KCl stimulation of 10 min. (L) The inhibited insulin release in the Eno 1 KD cells after 50 mM KCl stimulation of 10 min or incubation with 16.7 mM glucose for 1 h. All experiments were repeated independently at least 3 times. Significance test performed with single factor ANOVA test. * p<0.05; **, p<0.01; ***, p<0.001.

To corroborate the effects of the granule proteins on the size of IGCs, we performed transmission electron microscopy (TEM) imaging in β cells and analyzed the core size under the silencing conditions (Figure 3E). In line with the results from dSTORM imaging, IGCs were enlarged after silencing Nucb2 and Eno1 (Figure 3F). Silencing the granular H^+^ transporters, either cytosolic V1 domain ATP6V1a, or transmembrane V0 domain ATP6V0a1 (Figure 3A and B), did not result in changes in sizes of IGCs, as detected by dSTORM imaging. Pharmacological inhibition of the H^+^ transporter by Bafilomycin A1 also failed to change the IGCs sizes even though it was concomitant with reducing the number of insulin granules (Figure S3C, D). However, the IGCs size was significantly reduced in ATP6V1a silenced cells by TEM imaging (Figure 3F). As TEM detects the whole core electron density rather than insulin molecule density, we surmise that the altered H^+^ content in granules changes the composition/conformation of other proteins without affecting the insulin cores.

To test whether these proteins involved in regulating IGCs size participate in insulin secretory function, we firstly measured proinsulin and insulin content in the Nucb 2 or Eno1 silenced INS-1 cells (Figure 3G-I). In contrast to the increased size of IGCs in Nucb2-silenced cells (Figure 3A), the expression of proinsulin and insulin is rather decreased in the knock-down cells. However, the ratio of proinsulin/insulin indicated an increase of the proinsulin protein (Figure 3I and J). The amount of activity (50 mM KCl) but not 20 mM glucose-stimulated insulin secretion increased markedly in the Nucb2-silenced cells (Figure 3K and Figure S3E), consistent with the increased size of the IGCs in the cells. Meanwhile, silencing Eno-1 which polarized IGCs (Figure 3D) enhanced proinsulin and insulin expression, but KCl or glucose-stimulated insulin secretion was strongly inhibited (Figure 3L). These data demonstrate that Nucb2 and Eno-1 regulate insulin secretion by participating in the control of IGC’s size and shape rather than control of insulin gene expression.

### The size of IGCs in human primary β-cells is reduced upon glucose stimulation

To estimate the dynamic changes of exocytotic IGCs upon stimulation, we performed dSTORM imaging under TIRF mode on isolated primary human β-cells after incubation with 2.8 mM or 16.7 mM glucose (Figure 4A, B and Figure S4A). We found that stimulation with high glucose decreased the size of the insulin cores from 121.78±3.5 nm to 103.26±2.2 nm (Figure 4C), while the size measured by conventional microscopy remained unchanged (Figure 4D). Among the insulin cores, the population of larger sizes (>80 nm) decreased accordingly in response to the stimulation, while the population of smaller sizes (≤80 nm) increased (Figure 4E). By co-staining with VAMP2, we further confirmed that the stimulation-induced reduction of the larger cores locate preferentially in the exocytotic granules but not in non-exocytotic granules (Figure 4F and G). Accordingly, the size distribution of VAMP2 was also decreased after glucose stimulation (Figure 4H). However, the number of VAMP2-positive IGCs at the plasma membrane remained unchanged (Figure 4I). These results indicate that the stimulation-induced reduction of larger cores was not due to complete fusion but rather to incomplete granule fusion (kiss-and-run)(MacDonald et al., 2006).

**Figure 4.**
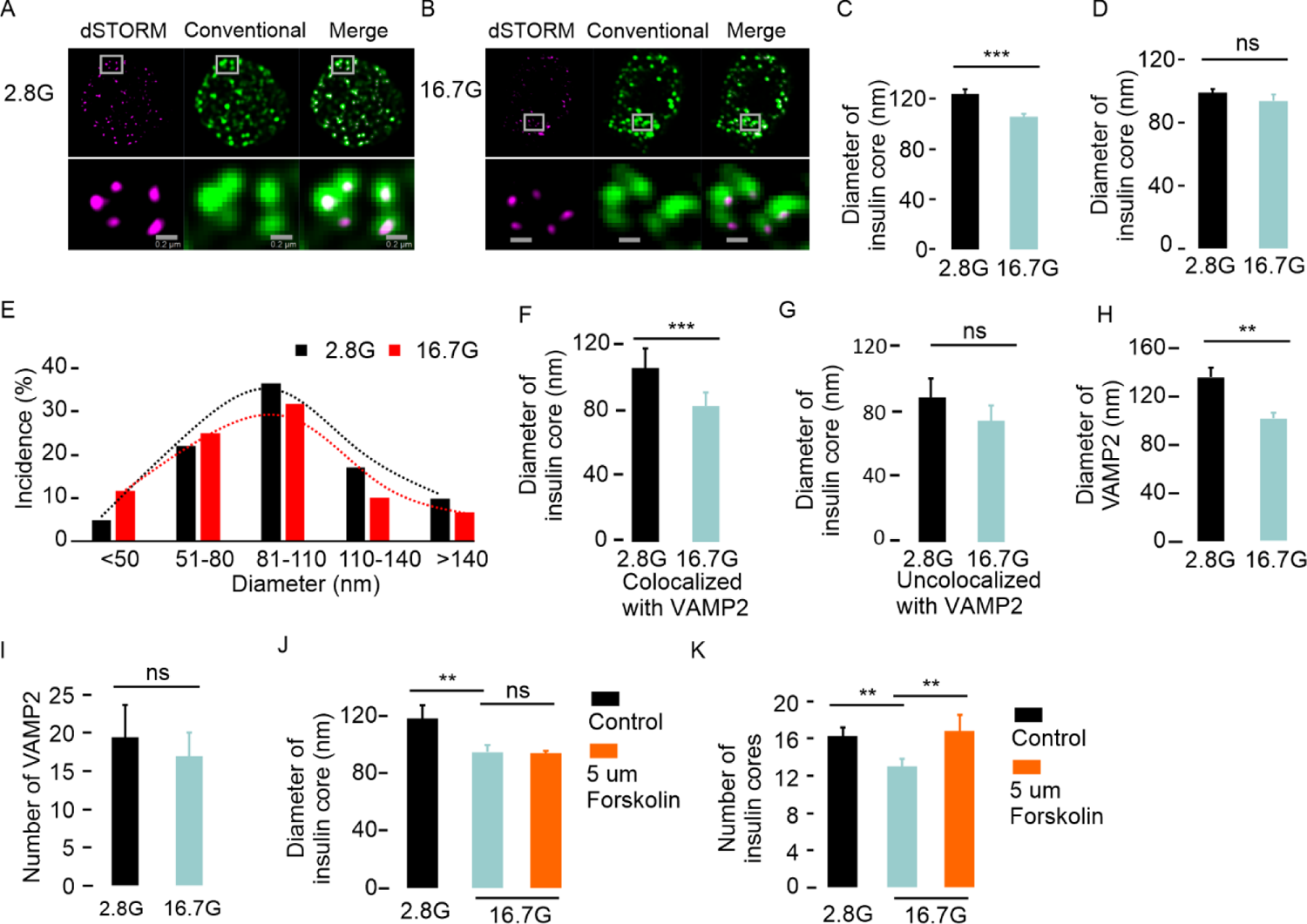
Reduction of insulin granule core size after stimulation of primary human β-cells. (A) Representative images of insulin cores under TIRF mode after 1-h incubation in 2.8 mM or (B) 16.7 mM glucose, respectively. (C) Decrease of the diameters after stimulation with 16.7 mM glucose under conditions as in (A), but no significant change was visualized by conventional imaging (D). (E) the distribution curve of the insulin cores indicated that the reduction mainly results from the population of larger size. (F) The size of insulin cores decreased in the granules that colocalized with the exocytotic marker VAMP2, but not in the granules without VAMP2 (G). (H) VAMP2 size reduced after 16.7 mM glucose stimulation. (I) VAMP2 numbers on the surface detected by TIRF mode remained non-significantly changed after 16.7 mM glucose stimulation. (J) Detection of insulin cores under stimulation with 5 uM forskolin plus 16.7 mM glucose stimulation for 30 min. (K) Number of surface insulin cores detected under TIRF mode after 1 h treatment with 5 uM forskolin and 16.7 mM glucose. Significance test performed with single factor ANOVA test. * p<0.05; **, p<0.01; ***, p<0.001, ns, no significance.

We next measured the number of insulin molecules in IGCs by accounting active blinking photons, which indirectly indicates the binding of dyes to insulin molecules. Remarkably, the photon number within the insulin core was significantly lower after glucose stimulation (Figure S4B and C), showing that part of the insulin core was released. Moreover, we treated the cells with 5 uM forskolin promoting incomplete fusion(MacDonald *et al*., 2006). Surprisingly, forskolin blocked the size reduction induced by glucose stimulation, confirming that the size reduction depended on the incomplete fusion (Figure 4J). In addition, the number of juxta-membrane insulin cores increased after forskolin treatment (Figure 4K), providing evidence that incomplete fusion events occurred during the stimulation. Taken together, these results demonstrate that the reduction of the core size by glucose is due to the release of part of the insulin core through incomplete granule secretion.

### The releasable part of IGCs is secreted through incomplete fusion upon glucose stimulation

To corroborate whether the size change is caused by the incomplete fusion, we developed a new protocol, which employs the SD antibody as a probe incorporating into IGCs during the glucose-stimulated granule fusion (Figure 5A). The diameter of the fusion pore of incomplete fusion is 1.8-6.3 nm depending on the species (Hanna et al., 2009), allowing the passage of the SD antibody (∼1 nm). The INS-1 cells were treated with glucose from 5 min to 1 hour, followed by the measurement of incorporated SD-probes by dSTORM imaging. We found that a significant number of labelled IGCs were detected after 5 min stimulation (Figure 5B). Accordingly, the size of the IGCs (∼90 nm, Figure S5A) with incomplete fusion was smaller than integral IGCs (118±23 nm, Figure 1D). The number of labelled IGCs declined following the glucose stimulation, while the average IGCs size remained unchanged. We then checked which size of the IGCs population was altered during the one-hour stimulation. Notably, in this case, the increased population of small size IGCs (>50-70 nm) is reciprocal to the decreased large IGCs. This reflects the releasable parts of the IGCs after incomplete fusion, detected predominantly at 5-min after stimulation with glucose (Figure 5C). This population represents the immediate releasable pool of the first phase in the phasic secretion(Gandasi et al., 2017). Later during the secretion, the large size of labelled IGCs (>100 nm) indicated the small releasable part was increased at 60-min stimulation. Taken together, these results confirm a dynamic change of the releasable zone in IGCs that reflects the glucose stimulation.

**Figure 5.**
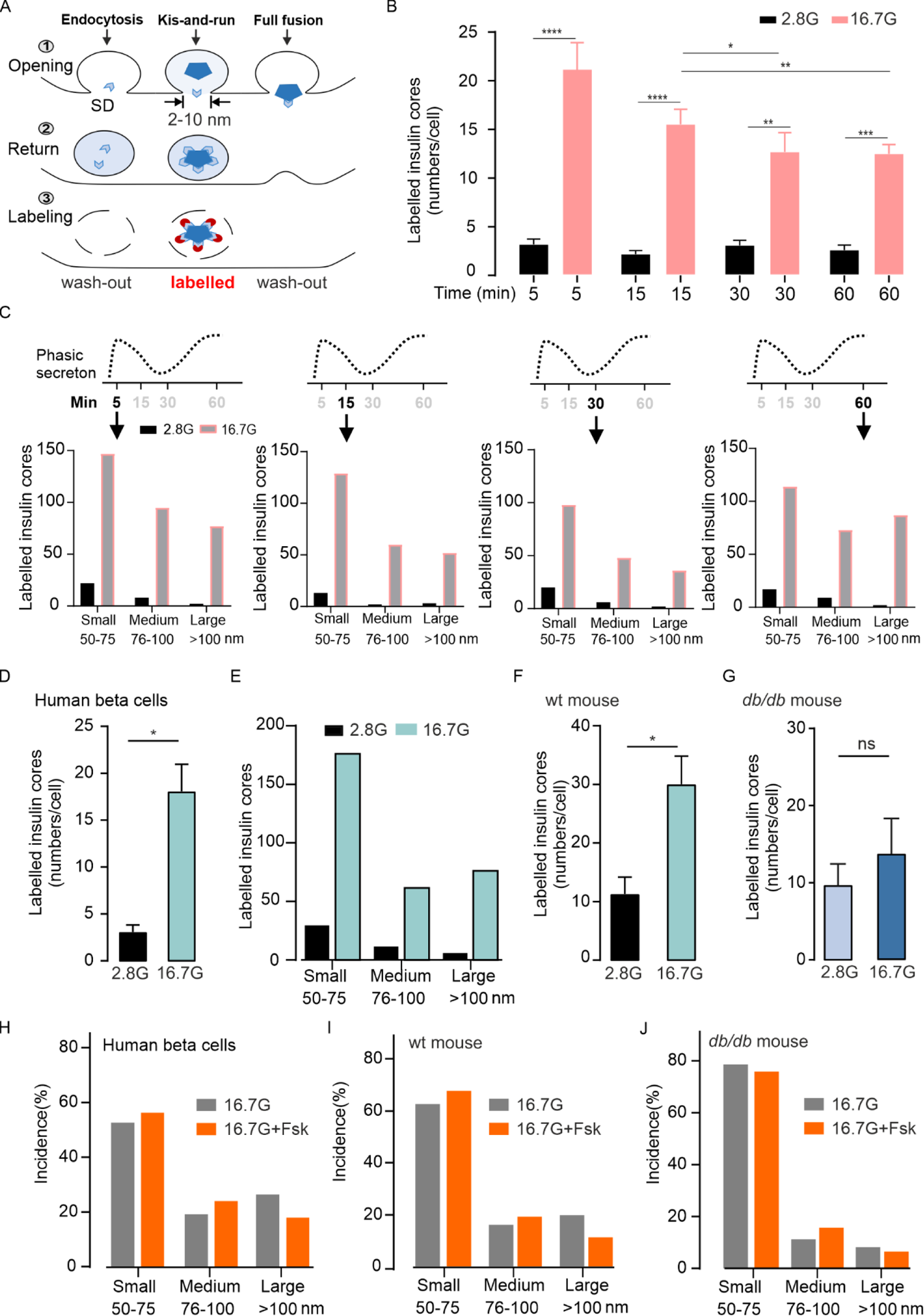
Partial IGCs are released through the incomplete fusion of insulin granules. (A) Schematic of labelling IGCs during incomplete fusion (Kiss-and-run). Step 1, SD antibodies were incubated with the cells during the stimulation. Step 2, SD-antibodies entered granules and bound to IGCs, in addition to some unspecific binding or endocytosis entry. Step 3, after standard staining protocol, only SD antibodies binding with IGCs remained. (B) The labelled insulin cores (number per cell) were measured in INS-1 832/13 cells after 2-h pre-incubation in 2.8 mM glucose then 5, 15, 30 and 60 min stimulation in 2,8 or 16,7 mM glucose with 1:200 insulin SD antibodies, Data presented as means ± SEM. (C) Distributions of labelled IGCs according to the IGCs sizes. Same conditions as B. (D) The labelled insulin cores were measured in human beta cells during 60 min stimulation. Data was collected from 3 donors. (E) The distribution of IGCs in human beta cells. (F) The labelled insulin cores measured in beta cells of wild type mouse and (G) db/db mouse for 60 min stimulation. The distribution of IGCs after 5 uM Forskolin treatment in beta cells of human (H), wild type mouse (I) and *db/db* mouse beta cells (J). Significance test performed with single factor ANOVA test. * p<0.05; **, p<0.01; ***, p<0.001; ns, no significance.

We next utilized the incomplete fusion protocol to assess whether this alteration of IGCs in the kiss-and-run model is a general phenomenon in healthy human beta cells as well as in primary mouse beta cells (Figure 5D and 5F). Similar distribution patterns of incomplete fusion events were observed in both cell types (Figure 5E, S5B). Of note, in the diabetic *db/db* mouse, the incomplete fusion events were not significantly changed after glucose stimulation, suggesting that the releasable zone in IGCs is defective in the diabetic mouse which indeed has impaired insulin secretion (Figure 5G) (Zhang *et al*., 2019). We further investigated the change of the releasable part in IGCs under the incomplete granule fusion by treatment with 5 uM forskolin, which promotes the fusion(Hanna *et al*., 2009). Forskolin treatment increased the population of small and medium sizes, but not the large size IGCs population in healthy human and mouse beta cells (Figure 5H and I). This confirms that a population of IGCs with a larger releasable zone was released upon forskolin treatment. Consistently, Forskolin did not affect the already decreased IGCs size in the *db/db* mouse (Figure 5J), indicating a defect in the releasable part in the IGCs.

### The size of IGCs is reduced in human diabetic β-cells

To further verify whether the defect of the releasable zone in IGCs also exists in another diabetic animal model, we next measured the core sizes in *Akita* (*Ins2^+/-^*) mice, which have a point mutation of the *ins2* gene, leading to incorrect folding and processing of proinsulin in the ß-cells, resulting in the development of severe hyperglycemia (Hong et al., 2007). Similarly, the size of the IGCs docked on the plasma membrane (under TIRF mode) was significantly reduced in cells from *Akita* mice when compared to non-diabetic control mice (Figure 6A). In contrast, the size of IGCs in cytosolic granules (under EPI mode) was not different (Figure 6B). Accordingly, photon numbers within the IGCs were also lower in cells from *Akita* mice (Figure S6A).

**Figure 6.**
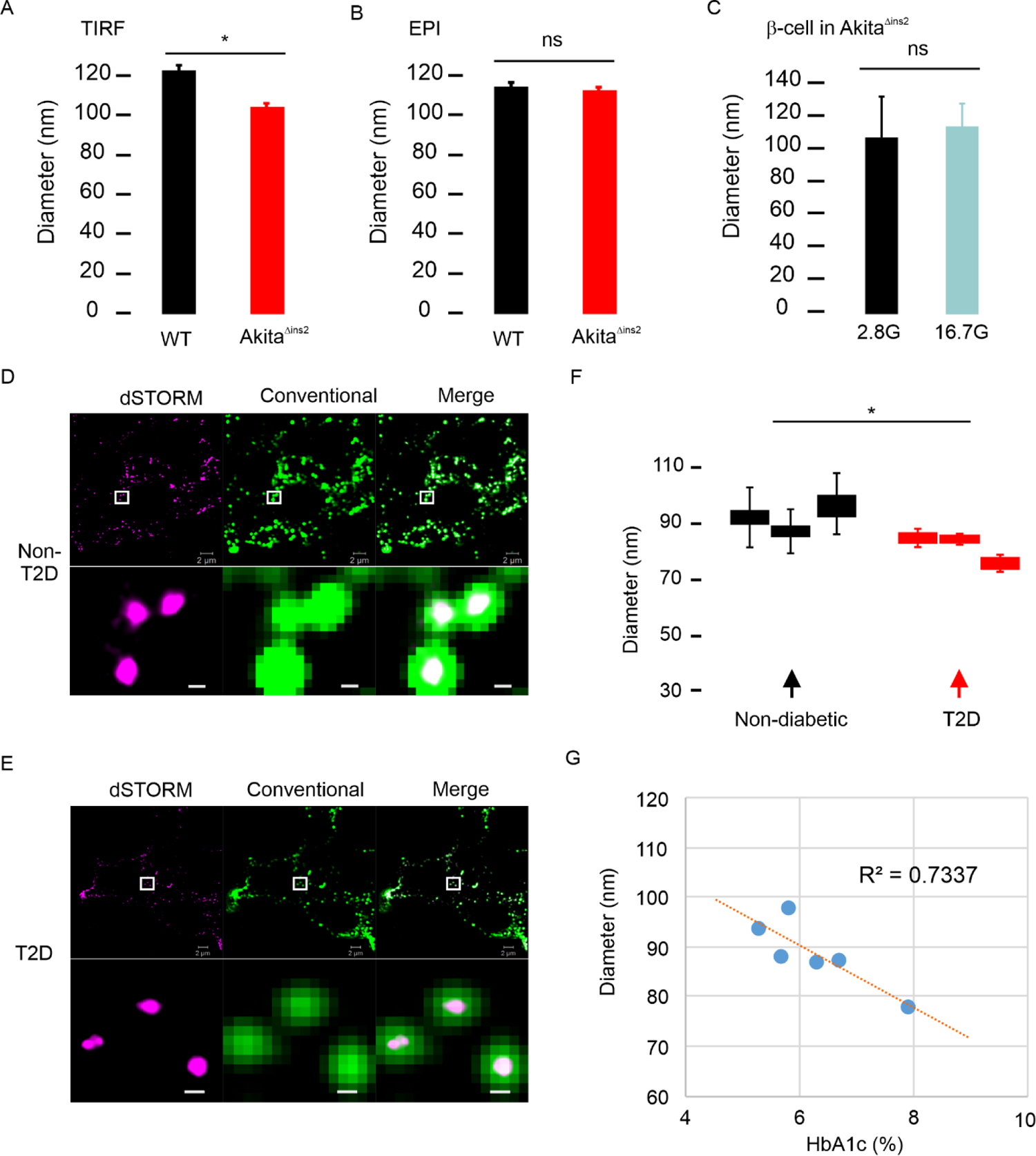
The insulin cores in β-cells from T2D donors are smaller than those of non-diabetic. (A) Insulin core size in granules under TIRF mode was reduced in the Akita^Δins2^ mice but remained unchanged under EPI mode (B). (C) Glucose stimulation was unable to further reduce the insulin cores in the Akita^Δins2^ mice. (D) Representative dSTORM images of insulin cores were acquired from the sections of the human pancreases from a healthy donor. The magnified images are from the squares as indicated in the respective upper images. (E) Same as in (D), but the images are acquired from a donor with type-2 diabetes. (F) The averages of average diameters of insulin cores were measured in healthy and diabetic donors. (G) the correlation between the blood HbA1c and the size of IGCs in the human donors. Significance test for the experiments performed with single factor ANOVA test. * p<0.05; ns, no significance.

**Figure 7.**
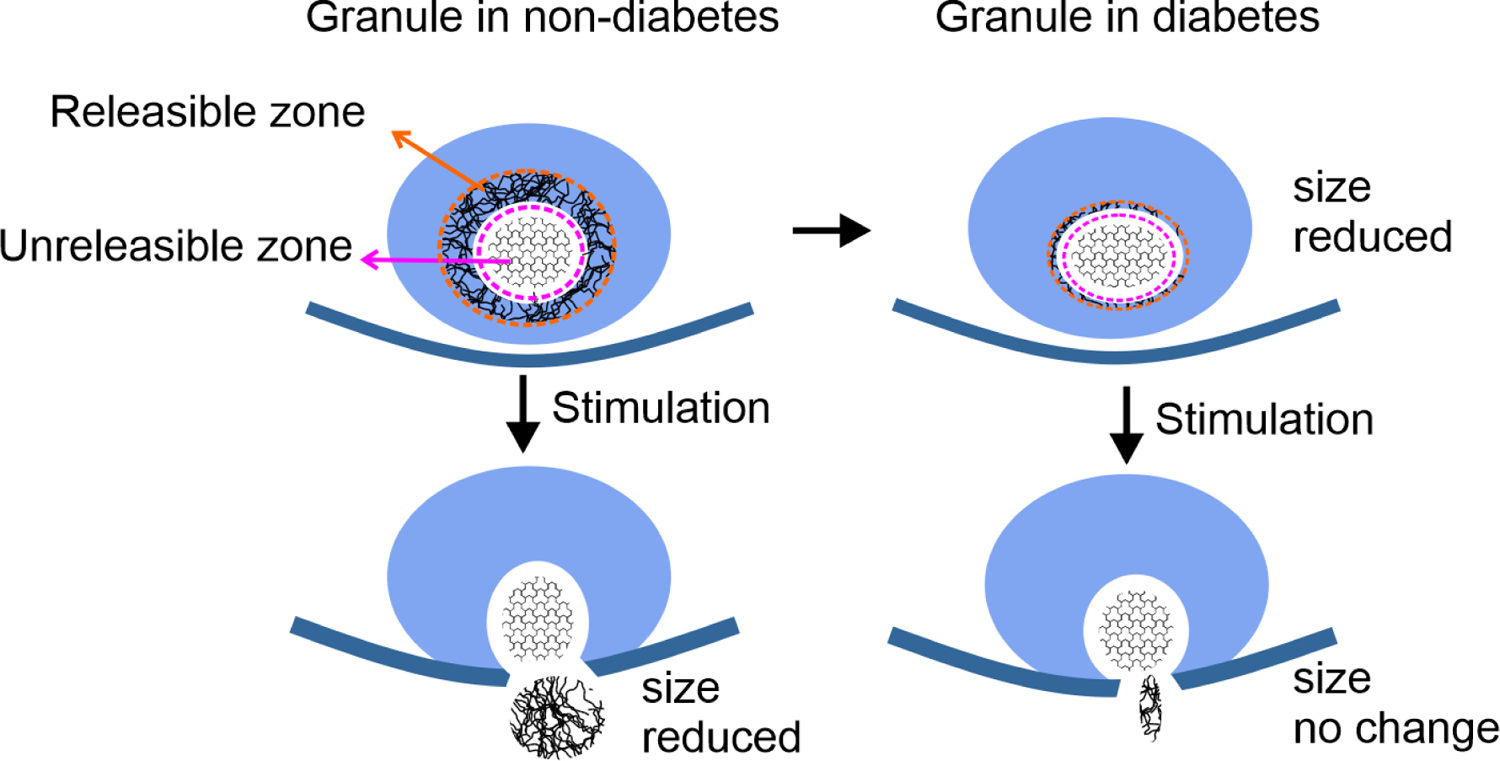
Schematic illustration of insulin core reduction in β-cells of T2D islets is associated with deficiency of glucose-stimulated insulin secretion. The insulin core is subdivided into the releasable zone and unreleasible zone. The releasable zone is secreted during the granule with incomplete fusion and resulted in reduction of core size. In the diabetic β-cell, the releasable zone is diminished, and therefore the size of the insulin cores remains unaltered after stimulation.

The size of IGCs that colocalized with Vamp2 was slightly smaller in *Akita* mice (Figure S6B). In addition, acute stimulation with 16.7 mM glucose of β-cells from *Akita* mice had no effect on the IGCs sizes (Figure 6C). These results showed that the releasable part of the insulin cores in the diabetic β-cells is diminished or deranged, further suggesting that reducing the releasable zone correlates with the inhibition of proinsulin cleavage and insulin secretion.

We next tested whether the reduction of the releasable part of the IGCs is present in human diabetic beta cells. To this end, we performed dSTORM imaging in intact paraffin-embedded human pancreases from diabetic and non-diabetic donors (Figure 6D and E). The sizes of IGCs in the three healthy donors were 91.6±11.1 nm, 88.5±9.4 nm, 97.2±13.4 nm, while in T2D donors, the IGCs size was significantly smaller with 86.2±4.4 nm, 85.7±1.8 nm, 76.5±3.8 nm, respectively (Figure 6F). Despite the large variability observed in the samples from healthy donors, a tendency for decreased photon numbers was clear (Figure S6C), indicating reduction of insulin molecules in the IGCs. The correlation analysis of IGCs size with glycated blood hemoglobin A1c (HbA1c) showed a strong negative correlation (r^2^=0.73) (Figure 6G). These findings indicate that reduction of the releasable zone in IGCs may result in defective exocytosis and contribute to the development of diabetes.

## Discussion

Using super-resolution dSTORM imaging combined with single-domain (SD) antibody, we provide evidence that the structure of the IGCs is dynamically changed in response to extracellular stimuli, secreting their releasable zone during the stimulation. Our unprecedented protocol enabled us to specifically characterize the ultrastructure of IGCs of the *in situ* granules and to describe a defect of the structure within IGCs in diabetic pancreatic β-cells.

dSTORM imaging provides considerable advantages to reveal the unknown ultrastructure of insulin granules at the subcellular level. However, this application meets practical challenges to reach the super-resolution due to the limitations such as the size of antibodies. The total length of the probe is usually larger than 20 nm, exceeding ten times the size of SD antibody (1-2 nm). This feature makes SD antibody particularly suitable for high-accurate imaging compared to conventional IgG antibody (Figure 1A-E). Besides the resolution, the size heterogeneity of IGCs in beta cells is also a challenging obstacle for acquiring results. Using a combination of EPI and TIRF mode in dSTORM imaging, we can distinguish the variations of the insulin cores at different cellular locations (Figure 2C and S2A-C). For example, the large insulin cores appear predominantly in exocytotic granules at the cell surface (Figure 2E), relative to the smaller ones found in the cytosol. This suggests that the larger IGCs in the exocytotic granule could be implicated in insulin secretion. However, while many factors controlling granule exocytosis have been defined, such as granule size, location, age and movement dynamics(Duncan et al., 2003; Eliasson *et al*., 2008; Zhang *et al*., 2019), the role of the size or structure of IGCs have not been established. This is mainly due to the lack of specific tools such as dSTORM imaging.

According to the models of granule exocytosis, the IGCs changes during glucose-stimulated insulin secretion also can be categorized as 1) complete fusion of the large-size insulin core (Tsuboi et al., 2006), 2) incomplete fusion of part of the cores (MacDonald *et al*., 2006). Our current results support the latter since we have observed that upon acute stimulation by 16.7 mM glucose, most of the IGCs size reduction is due to the secretion of its releasable part both in the INS-1 cells and human β-cells (Figure 4J and K, Figure 5). We hypothesized that the insulin core exists as distinct parts: a releasable zone, secreted after stimulation and an unreleasable zone retained after stimulation. The releasable zone is defective and shrunk in diabetic β-cells, and the full size of IGCs remains unchanged after glucose stimulation (Figure 6H).

It was long thought that the secretory capacity is only dependent on the secretory machinery formed by the SNARE complex and associated elements(Rorsman and Renstrom, 2003; Sudhof and Rothman, 2009). This view neglected the importance of the granule content, which defines the secretory volume depending on its physical size, releasable part and composition. Especially in the cases of incomplete fusion, it is critical that the structure of the IGC is capable of rapidly transforming and releasing part of the core through the fusion pore, generally sized 2-10 nm in diameter within seconds in the plasma membrane(Hanna *et al*., 2009). Whether the core in the docked granule at the plasma membrane merely secrete its releasable part, or randomly release any part of the core was unclear. By stimulation with 16.7 mM glucose, we observed that the IGC size is reduced accordingly and acts in concert with the exocytotic machinery to fulfil the demand of secretion (Figure 5B). However, the molecular mechanism underlying the regulation of the conversion between the two IGC zones and their structural features need to be further investigated. For example, how the single IGC splits during incomplete fusion and what signal pathways regulate the separation, are open questions for understanding the mechanism. Probing the two parts and tracing their dynamics in response to the stimulation are likely to yield such information. In addition, investigation of dynamic changes of the granule compositions within IGCs by super-resolution imaging will be a new approach for the elucidation of the maturation and secretion competence of the insulin granule.

Besides the physical structure, the chemical composition in the IGCs has also recently been linked to insulin secretion. According to proteomics studies, the insulin secretory granule contains less than 150 proteins (Brunner *et al*., 2007; Schvartz et al., 2012). The population of granular membrane proteins such as VAMP2 are involved in the regulation of granule intracellular trafficking and release by interaction with other signal molecules. The other major part of proteins participates in granule maturation, primarily regulating granular pH, ion concentration and enzyme catalytic activities(Schvartz *et al*., 2012). It is essential to understand how the granular proteins contribute to regulate the IGC structure and further affect insulin secretion (Figure 3). For example, silencing the granular protein Nucb2 (nucleobindin 2) decreases the amount of proinsulin/insulin (Figure 3H), though the stimulated insulin secretion is rather increased (Figure 3K). This is likely due to the enlargement of the insulin core size with larger releasable zone (Figure 3A). In contrast, knock-down of Eno-1 changes the IGCs shape from polarized state to rounder (Figure 3D), which is unfavorable for secretion of the releasable part of IGCs through incomplete fusion. The consequence is impaired glucose-stimulated insulin secretion (Figure 3L), while the total insulin amount is increased (Figure 3H). These results indicate that granular proteins appear to participate in the regulation of the insulin secretory process by changing the physical shape of the IGCs. However, the underlying mechanism connecting insulin core structure with secretion remains to be clarified. Future studies will define how other granular proteins and granular environmental factors such as pH, control the exocytotic process. Granular pH has been reported to be involved in the aggregation of granule content as well as conversion of proinsulin to insulin (Colomer et al., 1996). However, we were unable to observe significant changes of insulin core size by inhibition of the granular H^+^ ATP pump either by silencing ATP6V0a1, ATP6V1a, or the pharmacological blocker Bafilomycin A. These results suggest an alternative way to compensate for the deficiency of granular H^+^ and to reduce the risk of variation of intra-granular pH. Therefore, development of a new protocol to directly visualize the maturation of the insulin granule will enable us to provide more details about the insulin core structure and insulin maturation.

In summary, using a super-resolution dSTORM imaging combined with SD antibody, we characterized the structure changes of IGCs of insulin granule during glucose stimulation in pancreatic beta cells (Figure 6H). IGCs present in exocytotic granules appeared larger in size than cytosolic IGCs. The insulin granule proteins such as Nucb2 and Eno-1 participate in maintaining the size and shape of the IGCs, respectively. Glucose stimulation resulted in reduced size of the exocytotic IGCs by releasing their peripheral part “releasable zone”. This release mainly occurred in the granules displaying incomplete fusion during glucose stimulation. In the diabetic *db/db* and *Akita* mice as well as in type-2 diabetes donors, the size of the IGCs was significantly reduced. Moreover, the IGCs size was negatively correlated with the blood HbA1c concentration in the human donors. These findings overall highlight a new strategy to regulate the stimulated secretion by targeting of the structure of exocytotic IGCs in insulin granules and may be extended to other hormone-containing granules as well as neural cells.

## Methods

### Antibodies

Anti-insulin VHH Single Domain Antibody tagged with biotin was purchased from Creative Biolabs (Cat# NAB-1554-VHH); guinea pig primary IgG anti-insulin antibody from EuroDiagnostica (B65-1); and mouse monoclonal anti-VAMP2 antibody from SYSY. The secondary anti-guinea pig and anti-mouse antibodies conjugated with fluorescence dyes Alexa 647, Alexa 546 and Alexa 488 were from donkey serum (Jackson Immuno Research).

### Animals

All animal protocols in this study were performed in accordance with the Malmö/Lund Animal Care and Use Committee (5.2.18-10992/18) and abided by the Guide for the Care and Use of Laboratory Animals published by the Directive 2010/63/EU of the European Parliament.

Adult heterozygous diabetic Akita (*Ins2^+/−^*) mice (stock number 003548, C57BL/6J background) were obtained from The Jackson Laboratory and bred at our animal facility to generate Akita and non-diabetic wild-type littermates. Animals had free access to tap water and were fed normal chow diet. Blood glucose (CONTOUR® meter; Bayer) and body weight were monitored regularly after weaning at 4 weeks of age. Only male mice were used because of the limited hyperglycemia observed in Akita female mice. Akita mice develop pronounced and sustained hyperglycemia from the 4^th^ postnatal week. Mice were euthanized by cervical dislocation after anesthesia with 3% isoflurane in oxygen (2 L/min) from 18 to 24 weeks of age, at the point the diabetes phenotype was confirmed by blood glucose >15 mmol/l in a drop of blood from a tail puncture.

Isolation of the mouse pancreatic islets was performed by retrograde injection of a collagenase solution via the pancreatic duct and the islets were then hand-picked in Hank’s buffer (Sigma-Aldrich) with 1 mg/ml BSA under a stereomicroscope at room temperature. The 60 islets were grouped and dispersed into single cells with calcium-free Hanks’ buffer and seeded on a 12-cm glass-bottom dish. Before staining, the cells were incubated overnight in a humidified atmosphere in RPMI medium supplemented with 5 mM glucose, 10% (vol/vol) fetal bovine serum, 100 IU ml^−1^ penicillin, 100 μg ml^−1^ streptomycin.

### Cell culture

INS-1 832/13 cells (kindly donated by Dr. C. B. Newgaard, Duke University, USA) were produced from rats and genetically inserted human insulin gene and secreting human insulin (Hohmeier et al., 2000). Cells from passage 60-70 were used and secretion was assayed regularly. Only cells with robust secretion (>10-fold stimulation by glucose) were selected for further staining. Before staining, INS-1 832/13 cells were cultured in RPMI-1640 (Gibco BRL, NY) with 11.2 mmol/l glucose and 2 mmol/l l-glutamine. The medium was supplemented with 10% fetal bovine serum (Sigma, USA), 1 mmol/l pyruvate, 10 mmol/l HEPES (Sigma, USA), 50 μmol/l 2-mercaptoethanol (Sigma, USA), 100 units/ml penicillin, and 100 μg/ml streptomycin (Thermo Fisher, USA). Cells were cultured on 100-mm Petri dishes (Nunc, USA) and incubated in 5% CO2/95% air at 37°C before transfer to the glass coverslips used for imaging.

### Isolation of human pancreatic islets

Human pancreatic islets were obtained through the Exodiab Human Tissue Laboratory and the Nordic Network for Clinical Islet Transplantation (Prof. Olle Korsgren, Uppsala University, Sweden) (Axelsson et al., 2017). Briefly, the human pancreas was perfused with ice-cold collagenase, cut into pieces and placed in a digestion chamber at 37 °C. The separation of endocrine and exocrine tissues was achieved by a continuous density gradient. Selected fractions were then centrifuged to enrich for islets. The purity of islets was measured by dithizone staining. From this suspension, islets were hand-picked under a dissecting microscope. The islets were incubated in culture medium containing 5.6 mM glucose in CMRL-1066 (INC Biomedicals) supplemented with 10 mM HEPES, 2 mM L-glutamine, 50 μg ml^−1^ gentamicin, 0.25 μg ml^−1^ Fungizone (GIBCO), 20 μg ml^−1^ ciprofloxacin (Bayer Healthcare), 10 mM nicotinamide and 10% human serum at 37 °C (95% O_2_ and 5% CO_2_) for 1–9 days before experiments. Donors were grouped according to HbA1c *i.e.* less than 6% (healthy), higher than 6.5% or history of diabetes (diabetic). There was no difference in islet purity between healthy and T2D donors.

### Isolation of primary pancreatic β-cells

The handpicked islets were collected and grouped in batches of 120 islets. The islets were then added into 1 ml calcium-free isolation buffer containing (in mM) 138 NaCl, 5.6 KCl, 1.2 MgCl_2_, 5 HEPES, 3 glucose, 1 EGTA and 1 mg/ml BSA (pH 7.4), followed by incubation to 37 °C water bath for 12 min. The β-cells are released from islets by mechanical disruption pipetting down-up ∼10 times. The disruption was stopped by addition of 9 ml culture medium. The cells were centrifuged by 1.500 rpm for 2 min and the supernatant discarded. The cells were re-suspended with 1 ml culture medium and gently pipetted. The resultant cell suspension was plated on glass-bottom dishes and maintained in culture medium for 24 h.

### Knock-down of targeted proteins by siRNA silencing

The small interfering RNA (siRNA) experiments of the targeted genes to CPE, ZntT8, Chromogranin A, Eno 1, ATP6V0a1, ATP6V1a, Cathepsin D, PDIA3, PCSK2, Nucleobindin 2 and siRNA control #2 were purchased from ThermoFisher Scientific, USA. Prior to transfection, INS-1 832/13 cells were seeded in six-well plates at a density of ∼5 x 10^5^ cells for 24 h and then transfected by Lipofectamine® RNAiMAX Transfection Reagent (Invitrogen, USA) according to the manufacturer’s instructions. After transfection for 48 h or 72 h, the cells were collected for mRNA expression measurement.

To detect the knockdown efficacy of each targeted gene, qPCR was used to measure their expression after siRNA treatment. Total RNA from the cells was extracted using RNAeasy (Qiagen, Hilden, Germany). Concentration and purity of the RNA was measured with a NanoDrop ND-1000 spectrophotometer (NanoDrop Technologies, DE) and RNA Quality Indicator (RQI) higher than 8.0 (Experion Automated Electrophoresis, Bio-Rad, USA) was considered as high-quality total RNA preparations. 0.5 μg RNA of each condition was used for cDNA synthesis with SuperScript KIT (Thermofisher Scientific, USA). For real-time PCR, a 10 μl of reaction mixture with 20 ng cDNA, 5 μl TaqMan Mastermix (Thermofisher Scientific, USA), and 100 nM TagMan gene expression assay (at a dilution of 0.05) was run in a 7900HT Fast Real-Time System. The amplification was performed: 50 °C for 2 min, 95 °C for 10 min, 40 cycles of 95 °C for 15 sec, and 60 °C for 1 min. The mRNA expression was calculated relative to the housekeeping gene HPRT mRNA in the same sample.

### Stimulated insulin secretion in cultured cells

The INS-1 832/13 cells were seeded on the 24-well plate and washed twice with SAB buffer (pH 7.2) containing 114 mM NaCl, 4.7 mM KCl, 1.2 mM KH2PO4, 1.16 mM MgSO4, 25.5 mM NaHCO3, 2.6 mM CaCl2, 20 mM HEPES, 0.2% BSA before stimulation. For preincubation, SAB buffer containing 2.8 mM glucose was added to the cells for 2 h at 37 °C. For glucose-stimulated insulin secretion, the cells were then incubated with buffer containing 16.7 mM glucose for 1 h. For high K^+^ stimulated insulin secretion, the cells were incubated with buffer containing 50 mM KCl buffer containing for 10 min. Insulin concentration secreted from INS-1 832/13 cells was measured with rat insulin high range ELISA kit (Mercodia, Sweden) and normalized according to protein content per well. For protein and insulin content analysis, the cells were lysed with RIPA buffer (50 mM Tris HCl pH 8, 150 mM NaCl, 1% NP-40/Triton X, 0.1% SDS, 0.5% sodium deoxycholate, 2 mM EDTA and 50 mM NaF) after stimulation. The lysate was centrifuged at 10,000g for 5 min (4 °C). The total protein and insulin content was directly analysed or samples stored at −20 °C for analysis later. Total protein was measured with the Pierce BCA Protein Assay Kit (Thermo Scientific).

### Immunostaining of cells

INS-1 832/13 cells or primary human β-cells were seeded on the #1.5 (0.17 mm) glass-bottom dishes (WillCo Well, Germany) for 24 hours prior to staining. For the staining, the cells received 2% PFA in PBS buffer pH 7.4 (containing 137 mM NaCl, 2.7 mM KCl, 8 mM Na_2_HPO_4_, and 2 mM KH_2_PO_4_) for 10∼12 min to fix the cells and then the cells were permeabilized with Perm buffer (Thermo Fisher, USA) for 10 min. Unspecific binding was blocked by a buffer supplemented with 5% normal donkey serum in PBS. Guinea pig anti-insulin antibody (1:350), single domain anti-insulin antibody (1:200) or mouse anti-VAMP2 antibody (1:100) were diluted in the 5% donkey serum-containing PBS overnight. The samples were washed three times with a washing buffer (the blocking buffer plus 0.5% tween-20) for 10 min each time. For biotin-streptavidin binding, 1 μM streptavidin-Alexa 647 in blocking buffer was added to the cells and placed at room temperature for 30-40 minutes. For primary and secondary antibody binding, anti-Guinea pig or anti-mouse secondary antibodies conjugated with Alexa488 or Alexa 546 were added to the cells for 2 hours. After the reaction, the cells were washed three times with the washing buffer with tween-20, but without donkey serum. The samples were stored at 4°C for dSTORM imaging within 3-4 days.

### Immunostaining for pancreas sections

Formalin-fixed paraffin-embedded human T2D and control pancreatic sections with 5 um thickness were deparaffinised by multiple washing steps with washing solutions containing ethanol in concentrations from 99% to 50% and followed by heat-induced antigen retrieval using Retrievit 2 buffer (Biogenex). The tissues were re-fixed by methanol plus 10% H_2_O_2_ for 10 min and washed with TBST buffer three times and each time for 3 min. The unspecific binding was blocked for 20 min by blocking buffer (5% normal donkey serum in PBS). For immunohistochemistry, the cells were allowed to react with the single domain anti-insulin antibody (1:200) or the primary antibodies guinea pig anti-insulin (1:400). Dye labelling was done with 1 μM Streptavidin-Alexa 647 or the secondary antibody applied was Alexa 488-conjugated anti-guinea pig (1:400), respectively. The tissues were washed three times each time 10 min with washing buffer containing 0.5 tween-20. After washing the stained tissues were refrigerated (4°C) in a black box for dSTORM imaging within 3∼5 days.

### *In situ* detection of IGCs in granules during incomplete fusion

INS-1 832/13 cells, primary human β-cells or primary mouse β-cells were seeded on the #1.5 (0.17 mm) glass-bottom dishes (WillCo Well, Germany) for 24 hours prior to staining. The cells were then washed with 1 ml secretion assay buffer (SAB) (114 mM NaCl, 4.7 mM KCl, 1.2 mM KH2PO4, 1.16 mM MgSO4, 25.5 mM NaHCO3, 2.6 mM CaCl2, 20 mM HEPES (pH 7.3), 0.2% BSA) with 2.8 mM glucose twice, and preincubated in SAB for 2 h at 37°C. The cells were then stimulated during static incubation for 5 min, 15 min, 30 min or 1 h in 500 μl SAB containing 2.8 or 16.7 mM glucose with SD antibody (1:200). Then the cells were directly fixed by 2% PFA in PBS buffer pH 7.4 (containing 137 mM NaCl, 2.7 mM KCl, 8 mM Na2HPO4, and 2 mM KH2PO4) for 10 min and permeabilized with Perm buffer (Thermo Fisher, USA) for 15 min. Unspecific binding was blocked by a buffer supplemented with 5% normal donkey serum in PBS. For biotin-streptavidin binding, 1 µM Streptavidin-Alexa 647 in blocking buffer was added to the cells and placed at room temperature for 30-40 min. The cells were washed three times with 5% normal donkey serum in PBS. After the reaction, the cells were washed three times with the washing buffer with tween-20, but without donkey serum. The samples were stored at 4°C for dSTORM imaging within 3-4 days.

### Imaging buffers

The dSTORM imaging was performed in the imaging buffer described previously(Dempsey *et al*., 2011). Briefly, the imaging buffer contained 1X TN buffer (50 mM Tris, pH 8.0), 0.5 mg/ml glucose oxidase (Sigma, USA), 25 μg/ml catalase (Sigma, USA) and 25 mM MEA (cysteamine); 1 M MEA was stored at −20°C and used immediately after thawing. The buffer was prepared and stored at 4°C for use within 5 days. Before dSTORM imaging, 50 % (w/v) stock glucose was added to the buffer with a final 10% glucose in the imaging buffer. The imaging buffer was only used for the day with imaging experiments.

### Microscopy setup

dSTORM imaging was performed on the ELYRA P1 imaging system (Zeiss Germany). The system included an inverted microscope with 100X oil immerse objective lens with 1.46 NA and configured for additional ultra total internal reflection fluorescence (uTIRF). The samples were mounted in the ZEISS level adjustable insert holder and placed on the PIEZO stage with focus lock function. The focus plane and imaging field were observed and selected by transmitting beam or full-length UV lights. The fluorescence dyes were excited by the selected three laser lines, 488 nm, 543 nm and 633 nm. Accordingly, filter sets #4 for collection of the emission lights were chosen dependent on the fluorescence dyes. For Alexa 488, the 488 nm laser line was used for excitation and the emission light filtered BP 495-550; for Alexa 546, a 543 nm laser line was used for excitation and the emission light filtered BP 570-620; for Alexa 647, a 633 nm laser line was used for excitation and the emission light filtered LP 655. The images were acquired onto a 256 x 256 pixel frame of an electron-multiplying charge coupled device (EMCCD) camera (iXon DU897, Andor).

### Acquisition imaging at EPI model or TIRF model

For dSTORM imaging, an aliquot of 200 µl of the imaging buffer as described above was added to the glass-bottom chamber, which was mounted by a level auto-adjusted holder (Zeiss, Germany). Single channel imaging was performed with single laser line for the according dyes through a TIRF geometry. For EPI model, the focus position was set as above as 2.5 um to the plasma membrane and the position was locked by Piezo autofocus functions which allowed the 5 nm vibration. For the TIRF model, the focus position was on the plasma membrane within 150 nm distance to glass surface. Under constant illumination with 488 nm, 543nm or 642 nm laser light excitation in the imaging plane, the images, of which typically 20,000 frames were collected with exposure time of 18 ms, were acquired continuously until the blinking molecules of one dye observed were negligible. To generate the dSTORM images, the PALM processing function in the ZEN software was applied. First, the overlapping signals were discarded using a multi-emitter model for the whole image sequences. Second, to distinguish the real signal peak, the mask size was set to 7 pixels for anti-insulin single domain antibody and anti-insulin IgG antibody. For both antibodies anti-insulin IgG and SD, the ratio of signal/noise was set to 6. After the filtration, the localization of blinking molecules was collected by 2D Gaussian function using a theoretical point spread function (PSF). The distribution of localization precision was calculated according to ZEN software.

### Two-channel dSTORM imaging

The two laser lines, 488 nm and 633 nm were used to excite Alexa 488 and Alexa 647 which labelled VAMP2 and insulin, respectively. The samples were alternately illuminated by the 633 nm and 488 nm laser lines. The filter set #2 for collection of the two emission lights was used through MBS 405/488/642 and EF BP 420-480/BP 495-560/LP 650 (Zeiss, Germany), which allows fast dual-channel acquisition with the same reflector module. To avoid unspecific excitation, the sample was orderly illuminated with the 633 nm laser line for 500 image frames followed by 500 images using the 488 nm laser line on the regions of interest (ROI) with 256 x 256 pixel frame. Both channels were acquired by the EMCCD camera with frequency of 18 ms per image. The image sequences were collected and the dSTORM images were rendered by ZEN software.

### Images analysis and rendering

The image sequences were analysed using ZEN software (Zeiss, Germany). Lateral and axial drift during acquisition was corrected after reconstruction of dSTORM images using the Drift function of ZEN software. The drift function was tested by a fluorescent beads as fiducial marker before it was used to correct the images. After drift correction, the images were proceeded by grouping function. Finally, the present dSTORM images were corrected according to the following parameters: photon number, precision size, and first frame.

Unspecific signals resulting from cellular auto-fluorescence and nonspecific antibody binding, at times appeared on dSTORM images as low-density distribution. To remove the unspecific background, the molecules were filtered out if the number of molecules was less than 5 in an area of 100 nm perimeter.

### Measurement of insulin core sizes

All images were exported in the raw TIF format for the analysis. The pixel density of localization for insulin cores was determined by Matlab with additional functions. The insulin core signals were segmented by using the function 2-D segmentation app. The area, average diameter and perimeter of each core were calculated by Matlab Region Analyser app.

### Transmission Electron Microscopy

The knock-down and control cells were collected and prepared for electron microscopy for analysis of insulin granules as previously described (Olofsson et al., 2002). Briefly, the cells were fixed in 2.5% Glutaraldehyde in fresh Millionig’s buffer (1.88% NaH2PO4·H2O (Sigma-Aldrich), 0.43% NaOH, pH 7.2) and refrigerated for 2 h. The cells were then post-fixed in osmium tetroxide (1%) for 1 h after a wash by Millionig’s buffer. The fixed cells were dehydrated and embedded in AGAR 100 (Oxford Instruments Nordiska AB, Sweden) for the 100 nm ultrathin sections. The sections were placed on Cu-grids and images were acquired by a JEM electron microscope (JEOL-USA. Inc., USA). The diameter of cores in insulin granule was calculated using ImageJ 1.52 (NIH, Bethesda, MD, USA).

### Statistical analysis of data

All data were presented as means ± SEM for the indicated number of observations or presented by a representative result obtained from at least three different experiments. The significances between different conditions were analysed by single factor ANOVA test and a p value less than 0.05 was considered a significant difference.

## Acknowledgements

This project was funded by the Knut and Alice Wallenberg Foundation (2014-0074). Human islets were provided by the Nordic Network for Clinical Islet Transplantation and the EXODIAB Human Tissue Lab. We are grateful for the expert technical help of Anna-Maria Veljanovska Ramsay and valuable comments from Prof. Sara Linse (Lund University) and Prof. Patrik Rorsman (Oxford University). This work was also supported by the Swedish Foundation for Strategic Research (LUDC-IRC #15-0067) and by the Swedish Research Council through a Strategic Research Area grant (EXODIAB #2009-1039) and project grants to E.R, L.E., M.F.G. and E.Z. Science and Technology Guiding grant of Xiamen to NC. The work was also supported by grants from Region Skåne to E.R. and L.E., the Swedish Heart & Lund Foundation to M.F.G., the Albert Påhlsson Foundation, the Crafoord Foundation, and the Diabetes Wellness foundation, to E.Z.

## Author contributions

M.B., X.J., M.N, N.C., Y.Y., L.C. performed the experiments. U.K. analysed data. M.F.G. generated and phenotyped the Akita colony and contributed to the animal experiments. C.B.W., A.B. and L.E. participated in data interpretation and preparation of the manuscript. E.R. designed the study, analysed data and wrote the manuscript. E.Z. performed the experiments, designed the study, analysed data and wrote the manuscript. All authors reviewed and edited the manuscript and approved the final version of the manuscript.

## Competing interests

The authors declare no competing financial interests.

## Supplemental information

**Table S1.** Donor characteristics for the islets used in the experiments

|  | **Number** | **Age** | **BMI** | **HbA1c (%)** | **Stimulatory**<br>**index** | **Purity (%)** |
| --- | --- | --- | --- | --- | --- | --- |
| Non-T2D | 13 | 63.61±3.7 | 25.9±0.98 | 5.56±0.08 | 6.2±1.9 | 77.3±5.1 |
| T2D | 6 | 59.8±4.3 | 33.73±4.1 | 6.43±0.35 | 2.53±0.67 | 67.7±9.7 |

**Figure S1.**
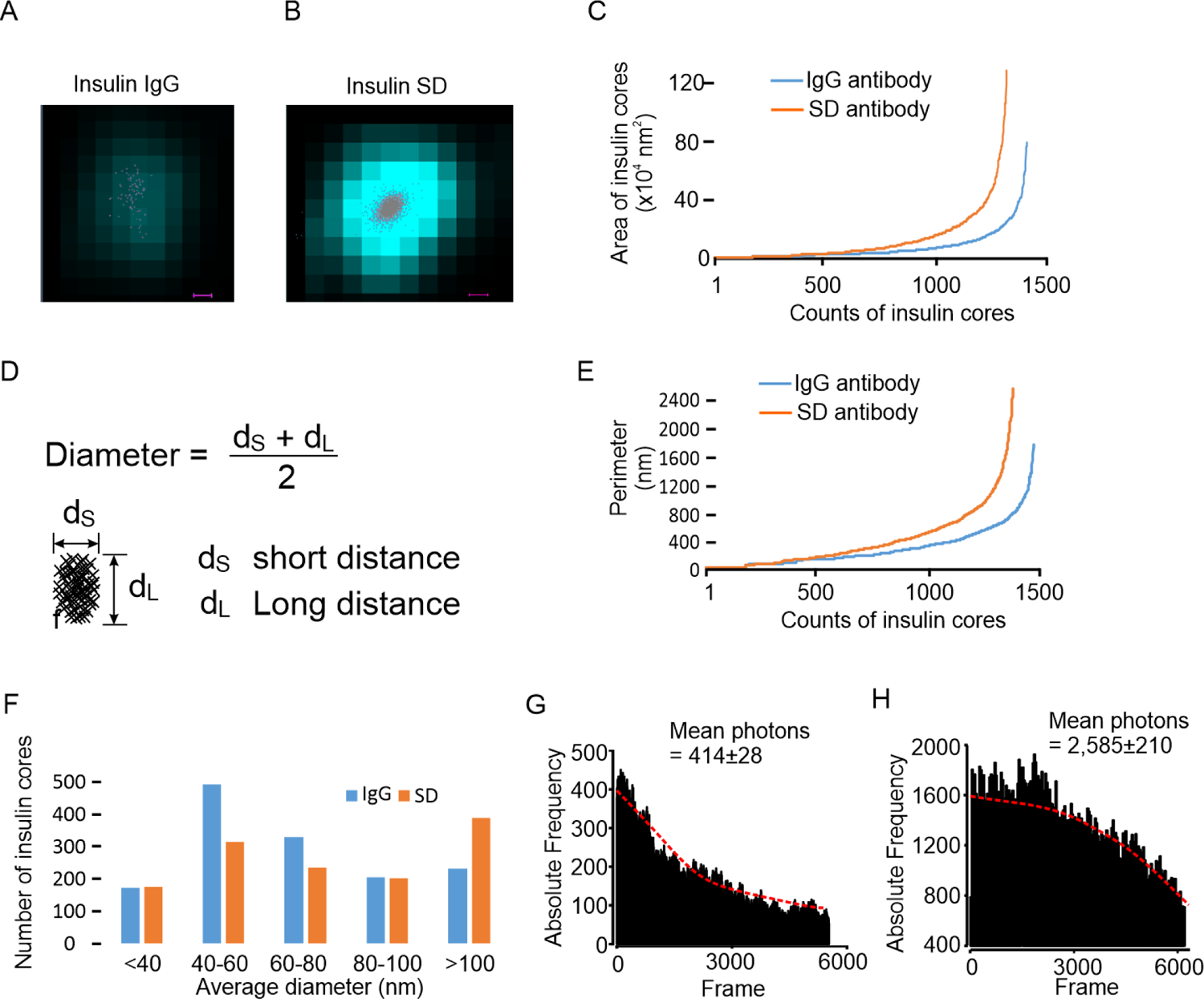
Higher accuracy of insulin core size when visualized by SD antibody. (A), distribution of molecular photons (grey spots) in a granular core with IgG antibody. The colour indicates the conventional image of the granule (Scale bar, 20 nm). (B), same as in a, but used SD antibody. (C) Area of insulin cores using IgG or SD antibodies detected by dSTORM imaging. The data were collected from 18 cells in each detection in three independent experiments. (D), the formula to calculate the diameter of insulin cores. (E) Same as in (C), but when measuring the perimeters. (F) The distribution of insulin cores detected by SD or IgG antibody-detected dSTORM imaging. Data are from 18 cells in three independent experiments. (G) The frequencies of photons per cell were measured using IgG insulin antibodies or SD insulin antibodies (H). Data presented are means ± SEM and collected from 18 cells in three independent experiments.

**Figure S2.**
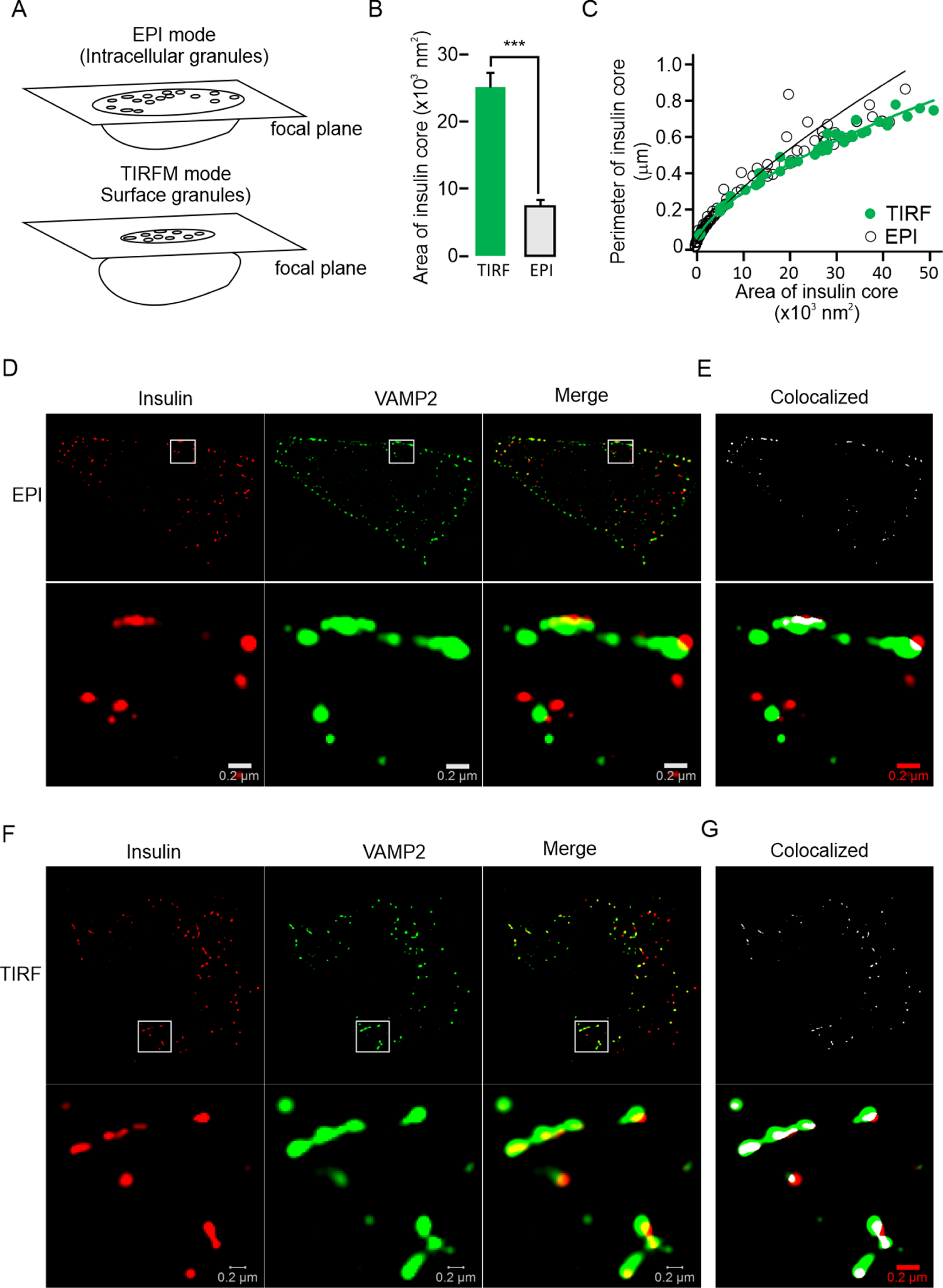
Large insulin cores locate in exocytotic granules. (A) illustrated the detection of EPI mode and TIRF mode in a single cell. (B) The average insulin core area was measured under EPI or TIRF mode. Significance test performed with single factor ANOVA test. ***p<0.001. (C) Comparison of correlations of perimeters and areas in EPI vs TIRF mode. The magenta hatched line indicates a positive linear correlation. Two-colour dSTORM images show the colocalization of insulin cores and VAMP2, an exocytotic large dense core insulin granule marker, in EPI (D) or TIRF mode (F). The processed images showed colocalization in white in EPI (E) or TIRF mode (G).

**Figure S3.**
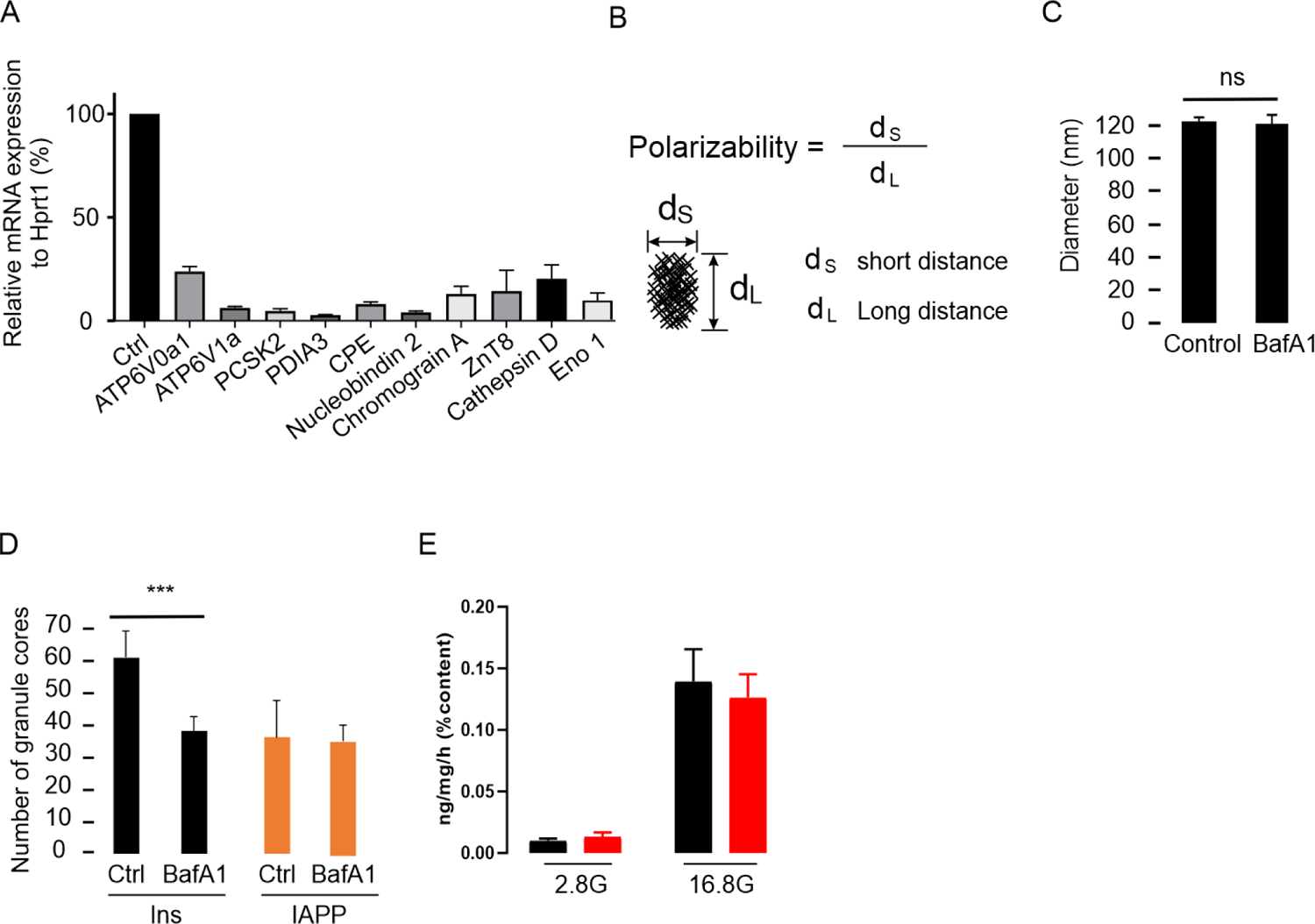
Detectable insulin molecules regulated after KD of insulin granular proteins. (A) Expression of the granular proteins after silencing by siRNA. (B) the formula used to calculate the rate of polarizability reflects the core shape from round (1) to long-stick state (0). (C) Insulin core size remained unchanged after 1 h of treatment with 0.5 μM bafilomycin A1, H^+^ ATPase inhibitor. (D) Surface insulin cores number detected by TIRF mode after treatment of bafilomycin A1. The IAPP size was also detected under the same conditions. Significance test for a-g performed with single factor ANOVA test. ***p<0.001 (E) Glucose stimulated insulin secretion has no significant change after KD of nucleobidin 2 even though it showed the reductive tendency. Significance test performed with single factor ANOVA test. * p<0.05; ns, no significance. All the experiments repeated at least three times independently.

**Figure S4.**
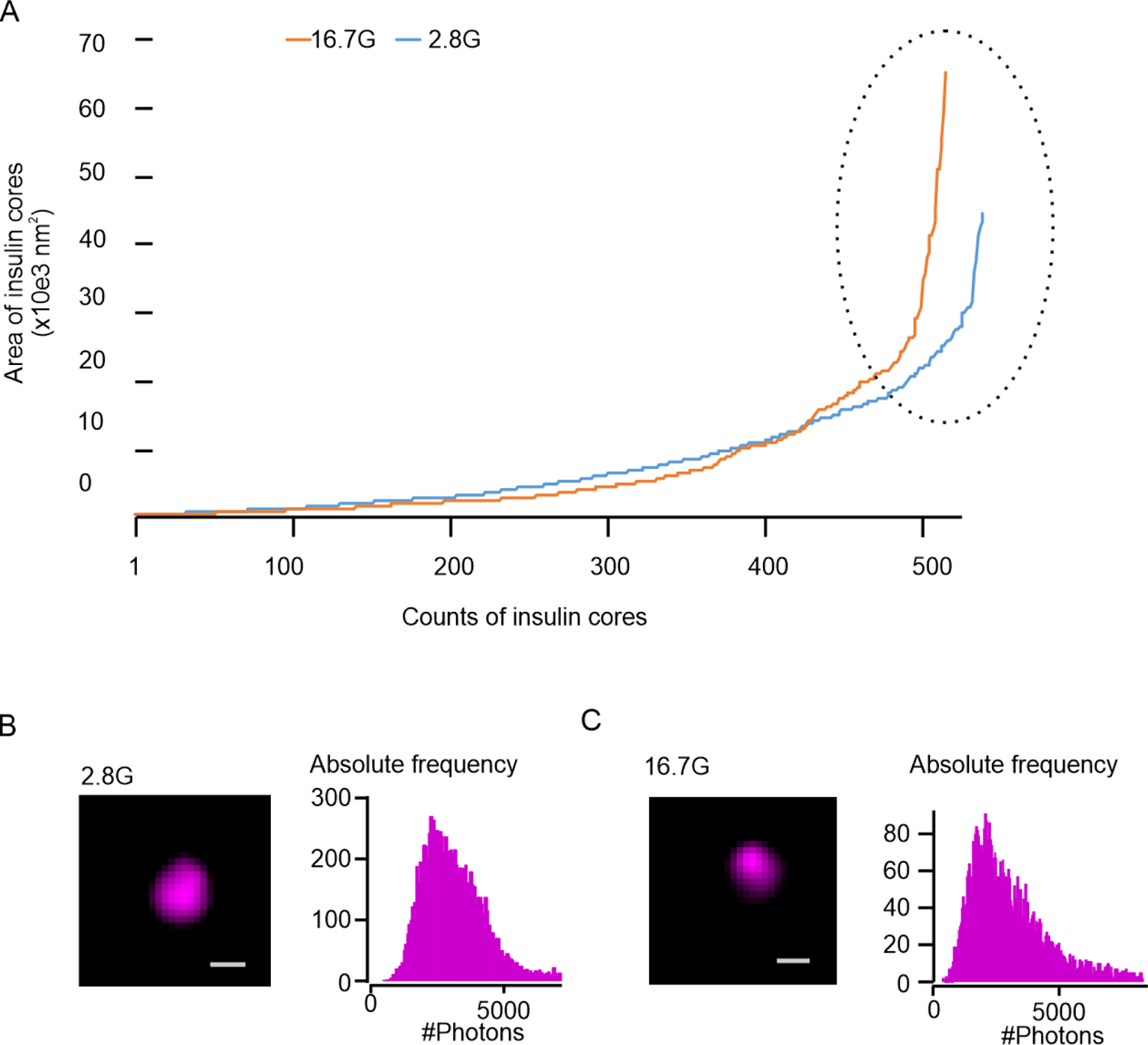
Glucose stimulation reduced insulin cores in primary human pancreatic beta cells. (A) Diameters of insulin core distributed from small to large with or without glucose stimulation. Notably, a smaller population (spot cycle) among large insulin cores showed under stimulation of glucose. (B) Representative image (left) and photon counts (right) in single insulin cores in cells incubated in 2.8 mM glucose. (C) same as in (B) but in glucose-stimulated (16.7 mM) cells.

**Figure S5.**
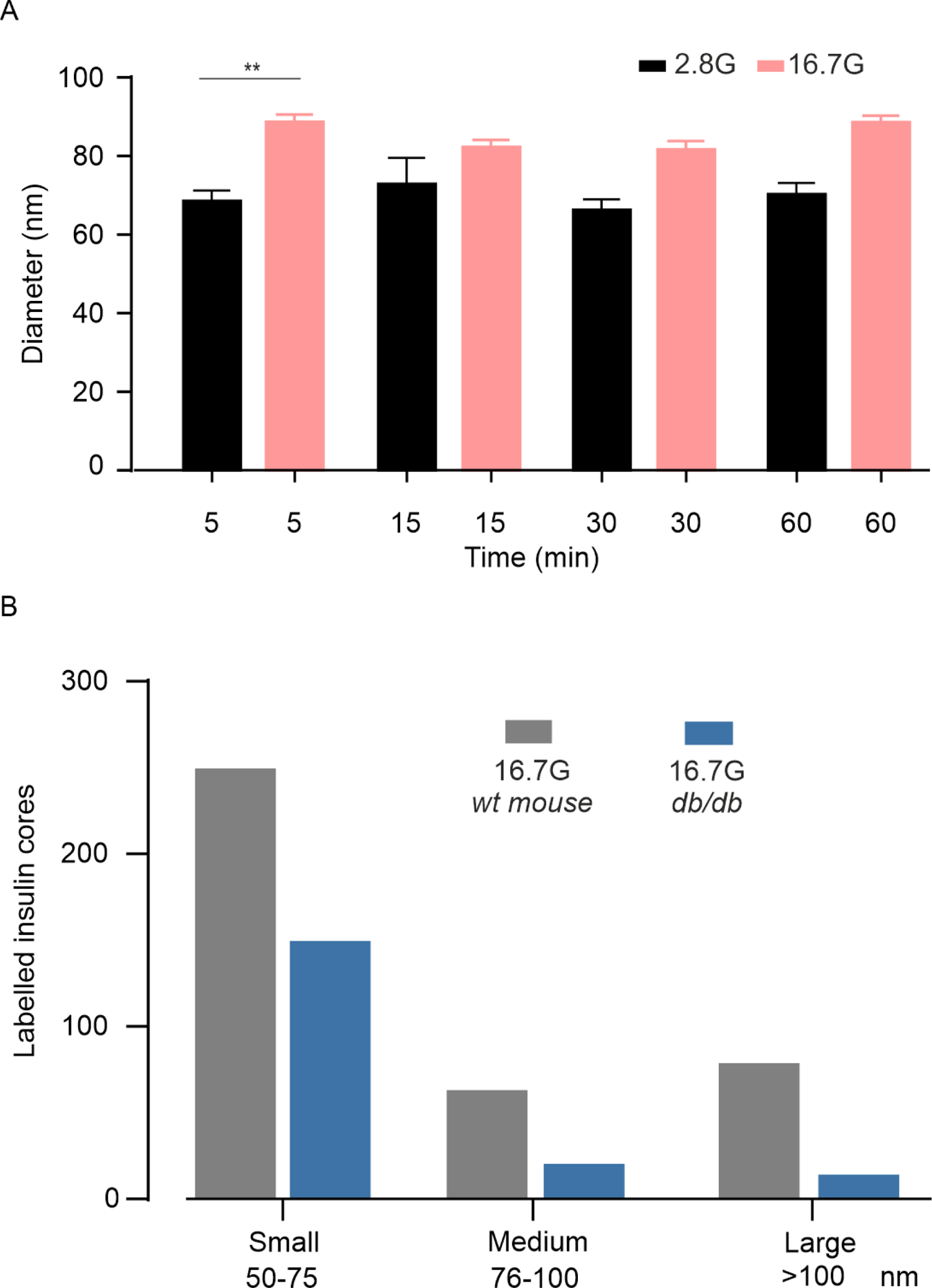
IGCs size changes in the granules during incomplete fusion. (A) The size of labelled insulin core in INS-1 832/13 cells after 2-h pre-incubation in 2.8 mM glucose then 5, 15, 30 and 60 min stimulation in 2,8 or 16,7 mM glucose with 1:200 insulin SD antibodies, Data presented as means ± SEM. (B) The distribution of IGCs in beta cells in the wild type (*wt*) and diabetic *db/db* mouse. Significance evaluated by single-factor ANOVA. **, p<0.01.

**Figure S6.**
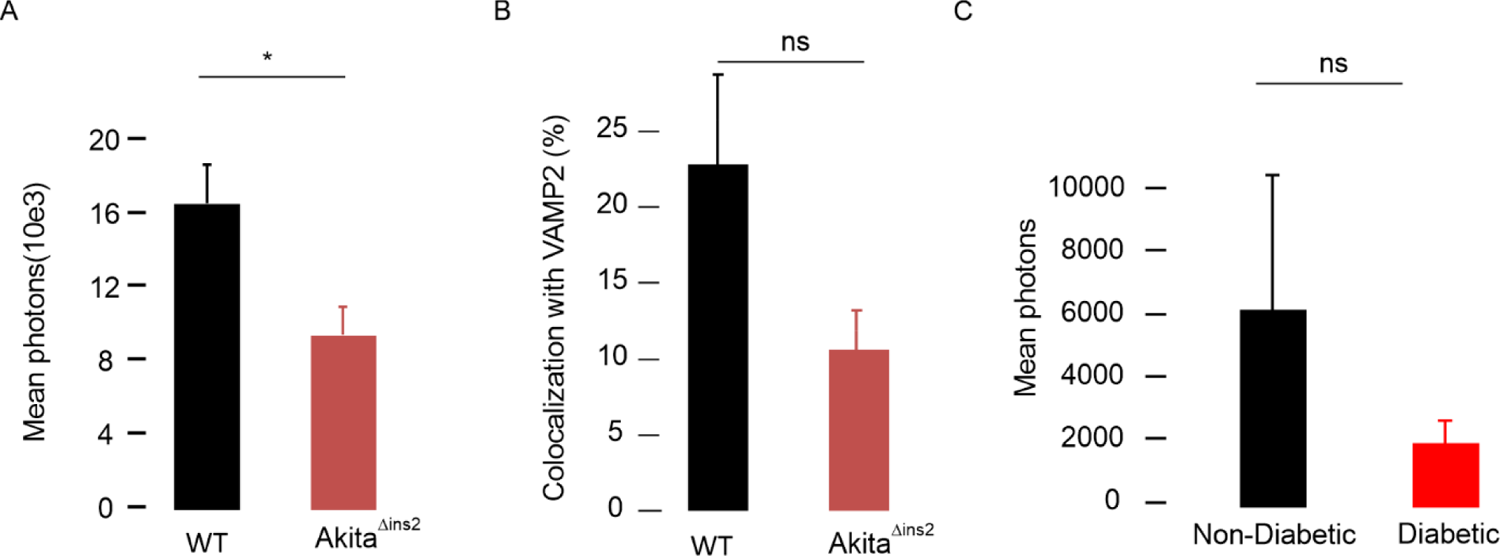
Reduction of IGCs size in the diabetic mouse and human beta cells. (A) Average mean photon counts per cell detected in Akita^Δins2^ mice. (B) The percentage of insulin core co-localized with Vamp2 in control and Akita^Δins2^ mice. (C) Average mean photon counts per cell in human b-cell. Data presented are means ± SEM and collected from 27 cells from each donor (three with diabetes and three without). Significance evaluated by single-factor ANOVA. *, p<0.05; ns, no significance.

## Notes

### Competing Interest Statement

The authors have declared no competing interest.

## References

1. Andersson, S.A., Olsson, A.H., Esguerra, J.L., Heimann, E., Ladenvall, C., Edlund, A., Salehi, A., Taneera, J., Degerman, E., Groop, L., et al. (2012). Reduced insulin secretion correlates with decreased expression of exocytotic genes in pancreatic islets from patients with type 2 diabetes. Mol Cell Endocrinol 364, 36–45. 10.1016/j.mce.2012.08.009.

2. Axelsson, A.S., Mahdi, T., Nenonen, H.A., Singh, T., Hanzelmann, S., Wendt, A., Bagge, A., Reinbothe, T.M., Millstein, J., Yang, X., et al. (2017). Sox5 regulates beta-cell phenotype and is reduced in type 2 diabetes. Nat Commun 8, 15652. 10.1038/ncomms15652.

3. Brunner, Y., Coute, Y., Iezzi, M., Foti, M., Fukuda, M., Hochstrasser, D.F., Wollheim, C.B., and Sanchez, J.C. (2007). Proteomics analysis of insulin secretory granules. Mol Cell Proteomics 6, 1007–1017. 10.1074/mcp.M600443-MCP200.

4. Colomer, V., Kicska, G.A., and Rindler, M.J. (1996). Secretory granule content proteins and the luminal domains of granule membrane proteins aggregate in vitro at mildly acidic pH. Journal of Biological Chemistry 271, 48–55. DOI 10.1074/jbc.271.1.48.

5. Cornu, M., Yang, J.Y., Jaccard, E., Poussin, C., Widmann, C., and Thorens, B. (2009). Glucagon-like peptide-1 protects beta-cells against apoptosis by increasing the activity of an IGF-2/IGF-1 receptor autocrine loop. Diabetes 58, 1816–1825. 10.2337/db09-0063.

6. Davidson, H.W., Rhodes, C.J., and Hutton, J.C. (1988). Intraorganellar calcium and pH control proinsulin cleavage in the pancreatic beta cell via two distinct site-specific endopeptidases. Nature 333, 93–96. 10.1038/333093a0.

7. Del Prato, S., Marchetti, P., and Bonadonna, R.C. (2002). Phasic insulin release and metabolic regulation in type 2 diabetes. Diabetes 51 *Suppl 1*, S109–116.

8. Dempsey, G.T., Vaughan, J.C., Chen, K.H., Bates, M., and Zhuang, X. (2011). Evaluation of fluorophores for optimal performance in localization-based super-resolution imaging. Nat Methods 8, 1027–1036. 10.1038/nmeth.1768.

9. Duncan, R.R., Greaves, J., Wiegand, U.K., Matskevich, I., Bodammer, G., Apps, D.K., Shipston, M.J., and Chow, R.H. (2003). Functional and spatial segregation of secretory vesicle pools according to vesicle age. Nature 422, 176–180. 10.1038/nature01389.

10. Eliasson, L., Abdulkader, F., Braun, M., Galvanovskis, J., Hoppa, M.B., and Rorsman, P. (2008). Novel aspects of the molecular mechanisms controlling insulin secretion. J Physiol 586, 3313–3324. 10.1113/jphysiol.2008.155317.

11. Gandasi, N.R., Yin, P., Omar-Hmeadi, M., Ottosson Laakso, E., Vikman, P., and Barg, S. (2018). Glucose-Dependent Granule Docking Limits Insulin Secretion and Is Decreased in Human Type 2 Diabetes. Cell Metab 27, 470–478 e474. 10.1016/j.cmet.2017.12.017.

12. Gandasi, N.R., Yin, P., Riz, M., Chibalina, M.V., Cortese, G., Lund, P.E., Matveev, V., Rorsman, P., Sherman, A., Pedersen, M.G., and Barg, S. (2017). Ca2+ channel clustering with insulin-containing granules is disturbed in type 2 diabetes. J Clin Invest 127, 2353–2364. 10.1172/JCI88491.

13. Hanna, S.T., Pigeau, G.M., Galvanovskis, J., Clark, A., Rorsman, P., and MacDonald, P.E. (2009). Kiss-and-run exocytosis and fusion pores of secretory vesicles in human beta-cells. Pflugers Arch 457, 1343–1350. 10.1007/s00424-008-0588-0.

14. Heilemann, M., van de Linde, S., Schuttpelz, M., Kasper, R., Seefeldt, B., Mukherjee, A., Tinnefeld, P., and Sauer, M. (2008). Subdiffraction-resolution fluorescence imaging with conventional fluorescent probes. Angew Chem Int Ed Engl 47, 6172–6176. 10.1002/anie.200802376.

15. Hickey, A.J., Bradley, J.W., Skea, G.L., Middleditch, M.J., Buchanan, C.M., Phillips, A.R., and Cooper, G.J. (2009). Proteins associated with immunopurified granules from a model pancreatic islet beta-cell system: proteomic snapshot of an endocrine secretory granule. J Proteome Res 8, 178–186. 10.1021/pr800675k.

16. Hoboth, P., Muller, A., Ivanova, A., Mziaut, H., Dehghany, J., Sonmez, A., Lachnit, M., Meyer-Hermann, M., Kalaidzidis, Y., and Solimena, M. (2015). Aged insulin granules display reduced microtubule-dependent mobility and are disposed within actin-positive multigranular bodies. Proc Natl Acad Sci U S A 112, E667–676. 10.1073/pnas.1409542112.

17. Hohmeier, H.E., Mulder, H., Chen, G., Henkel-Rieger, R., Prentki, M., and Newgard, C.B. (2000). Isolation of INS-1-derived cell lines with robust ATP-sensitive K+ channel-dependent and -independent glucose-stimulated insulin secretion. Diabetes 49, 424–430.

18. Hong, E.G., Jung, D.Y., Ko, H.J., Zhang, Z., Ma, Z., Jun, J.Y., Kim, J.H., Sumner, A.D., Vary, T.C., Gardner, T.W., et al. (2007). Nonobese, insulin-deficient Ins2Akita mice develop type 2 diabetes phenotypes including insulin resistance and cardiac remodeling. Am J Physiol Endocrinol Metab 293, E1687–1696. 10.1152/ajpendo.00256.2007.

19. Hu, Y.S., Cang, H., and Lillemeier, B.F. (2016). Superresolution imaging reveals nanometer- and micrometer-scale spatial distributions of T-cell receptors in lymph nodes. Proc Natl Acad Sci U S A 113, 7201–7206. 10.1073/pnas.1512331113.

20. Hutton, J.C., Penn, E.J., and Peshavaria, M. (1982). Isolation and characterisation of insulin secretory granules from a rat islet cell tumour. Diabetologia 23, 365–373.

21. Lemaire, K., Ravier, M.A., Schraenen, A., Creemers, J.W., Van de Plas, R., Granvik, M., Van Lommel, L., Waelkens, E., Chimienti, F., Rutter, G.A., et al. (2009). Insulin crystallization depends on zinc transporter ZnT8 expression, but is not required for normal glucose homeostasis in mice. Proc Natl Acad Sci U S A 106, 14872–14877. 10.1073/pnas.0906587106.

22. Ligthart, S., van Herpt, T.T., Leening, M.J., Kavousi, M., Hofman, A., Stricker, B.H., van Hoek, M., Sijbrands, E.J., Franco, O.H., and Dehghan, A. (2016). Lifetime risk of developing impaired glucose metabolism and eventual progression from prediabetes to type 2 diabetes: a prospective cohort study. Lancet Diabetes Endocrinol 4, 44–51. 10.1016/S2213-8587(15)00362-9.

23. Lyssenko, V., Jonsson, A., Almgren, P., Pulizzi, N., Isomaa, B., Tuomi, T., Berglund, G., Altshuler, D., Nilsson, P., and Groop, L. (2008). Clinical risk factors, DNA variants, and the development of type 2 diabetes. N Engl J Med 359, 2220-2232. 10.1056/NEJMoa0801869.

24. MacDonald, P.E., Braun, M., Galvanovskis, J., and Rorsman, P. (2006). Release of small transmitters through kiss-and-run fusion pores in rat pancreatic beta cells. Cell Metab 4, 283–290. 10.1016/j.cmet.2006.08.011.

25. Mengistu, M., Tang, A.H., Foulke, J.S., Jr., Blanpied, T.A., Gonzalez, M.W., Spouge, J.L., Gallo, R.C., Lewis, G.K., and DeVico, A.L. (2017). Patterns of conserved gp120 epitope presentation on attached HIV-1 virions. Proc Natl Acad Sci U S A 114, E9893–E9902. 10.1073/pnas.1705074114.

26. Nam, D., Mantell, J., Bull, D., Verkade, P., and Achim, A. (2014). A novel framework for segmentation of secretory granules in electron micrographs. Med Image Anal 18, 411–424. 10.1016/j.media.2013.12.008.

27. Ogawa, A., Harris, V., McCorkle, S.K., Unger, R.H., and Luskey, K.L. (1990). Amylin secretion from the rat pancreas and its selective loss after streptozotocin treatment. J Clin Invest 85, 973–976. 10.1172/JCI114528.

28. Olofsson, C.S., Gopel, S.O., Barg, S., Galvanovskis, J., Ma, X., Salehi, A., Rorsman, P., and Eliasson, L. (2002). Fast insulin secretion reflects exocytosis of docked granules in mouse pancreatic B-cells. Pflugers Arch 444, 43–51. 10.1007/s00424-002-0781-5.

29. Orci, L., Ravazzola, M., Amherdt, M., Madsen, O., Perrelet, A., Vassalli, J.D., and Anderson, R.G. (1986). Conversion of proinsulin to insulin occurs coordinately with acidification of maturing secretory vesicles. J Cell Biol 103, 2273–2281.

30. Ostenson, C.G., Gaisano, H., Sheu, L., Tibell, A., and Bartfai, T. (2006). Impaired gene and protein expression of exocytotic soluble N-ethylmaleimide attachment protein receptor complex proteins in pancreatic islets of type 2 diabetic patients. Diabetes 55, 435–440.

31. Regazzi, R., Sadoul, K., Meda, P., Kelly, R.B., Halban, P.A., and Wollheim, C.B. (1996). Mutational analysis of VAMP domains implicated in Ca2+-induced insulin exocytosis. EMBO J 15, 6951–6959.

32. Ries, J., Kaplan, C., Platonova, E., Eghlidi, H., and Ewers, H. (2012). A simple, versatile method for GFP-based super-resolution microscopy via nanobodies. Nat Methods 9, 582-584. 10.1038/nmeth.1991. Rorsman, P., and Renstrom, E. (2003). Insulin granule dynamics in pancreatic beta cells. Diabetologia 46, 1029-1045. 10.1007/s00125-003-1153-1.

33. Rothbauer, U., Zolghadr, K., Tillib, S., Nowak, D., Schermelleh, L., Gahl, A., Backmann, N., Conrath, K., Muyldermans, S., Cardoso, M.C., and Leonhardt, H. (2006). Targeting and tracing antigens in live cells with fluorescent nanobodies. Nat Methods 3, 887–889. 10.1038/nmeth953.

34. Rust, M.J., Bates, M., and Zhuang, X. (2006). Sub-diffraction-limit imaging by stochastic optical reconstruction microscopy (STORM). Nat Methods 3, 793–795. 10.1038/nmeth929.

35. Schvartz, D., Brunner, Y., Coute, Y., Foti, M., Wollheim, C.B., and Sanchez, J.C. (2012). Improved characterization of the insulin secretory granule proteomes. J Proteomics 75, 4620–4631. 10.1016/j.jprot.2012.04.023.

36. Suckale, J., and Solimena, M. (2010). The insulin secretory granule as a signaling hub. Trends Endocrinol Metab 21, 599–609. 10.1016/j.tem.2010.06.003.

37. Sudhof, T.C., and Rothman, J.E. (2009). Membrane fusion: grappling with SNARE and SM proteins. Science 323, 474–477. 10.1126/science.1161748.

38. Thurmond, D.C., and Gaisano, H.Y. (2020). Recent Insights into Beta-cell Exocytosis in Type 2 Diabetes. J Mol Biol 432, 1310–1325. 10.1016/j.jmb.2019.12.012.

39. Tsuboi, T., Ravier, M.A., Parton, L.E., and Rutter, G.A. (2006). Sustained exposure to high glucose concentrations modifies glucose signaling and the mechanics of secretory vesicle fusion in primary rat pancreatic beta-cells. Diabetes 55, 1057–1065.

40. Wang, Z., and Thurmond, D.C. (2009). Mechanisms of biphasic insulin-granule exocytosis - roles of the cytoskeleton, small GTPases and SNARE proteins. J Cell Sci 122, 893–903. 10.1242/jcs.034355.

41. Zhang, E., Mohammed Al-Amily, I., Mohammed, S., Luan, C., Asplund, O., Ahmed, M., Ye, Y., Ben-Hail, D., Soni, A., Vishnu, N., et al. (2019). Preserving Insulin Secretion in Diabetes by Inhibiting VDAC1 Overexpression and Surface Translocation in beta Cells. Cell Metab 29, 64–77 e66. 10.1016/j.cmet.2018.09.008.

